# The impact of homeostatic inhibitory plasticity in a generative biophysical model

**DOI:** 10.64898/2026.01.12.699008

**Authors:** Iván Mindlin, Carlos Coronel-Oliveros, Marc Llabrés, Jacobo D. Sitt, Rodrigo Cofré, Andrea Luppi, Thomas Andrillon, Yonatan Sanz Perl, Rubén Herzog

## Abstract

A main characteristic of biological systems is their capacity to dynamically adapt to environmental changes. In the brain, synaptic plasticity enables the strengthening or weakening of connections between neurons, allowing neural circuits to adapt based on experience, learning, and environmental changes. Yet, it is homeostatically regulated such that it avoids excessive proliferation of synaptic contacts. These mechanisms can be studied with large-scale models of brain activity. Here, we embed a biologically grounded inhibitory-homeostatic plasticity rule into the Dynamic Mean Field (DMF) model, creating a Homeostatic Dynamic Mean Field (HDMF) model that dynamically tunes local excitation–inhibition balance. Convergence of excitatory firing rates is reached by mapping a large range of coupling strength to parameters of inhibitory synapses. The HDMF reproduces statistical observables of brain activity as well as the original DMF, and can sustain neuromodulatory perturbations without overhead computations. The HDMF can generate unprecedented sleep-like slow-wave activity, which can also coexist with wake-like asynchronous dynamics, permitting to model dissociated states of consciousness such as parasomnias. Together, these results show that a single homeostatic rule broadens the stability and expressiveness of the DMF, providing a unified platform for studying how local adaptive processes shape the diverse global dynamics of the human brain.

## Introduction

Distinct brain dynamics underlie different states of arousal (Brown et al., 2011), consciousness (Luppi et al., 2019; Sanz Perl et al., 2021), cognition (Varela et al., 2001) , and health conditions (Dehaene et al., 2011). This varied landscape is grounded in the brain’s remarkable capacity to self-regulate the underlying physiological processes that induce them (Wen & Turrigiano, 2024). While some variables remain tightly controlled leading to stable functioning, others can shift over time, giving rise to the wide array of observable brain states (Foster & Scheinost, 2024). At the microscopic level, plasticity serves as an adaptive mechanism, continuously fine-tuning neural activity in response to a dynamic environment. However, synaptic changes are tightly regulated by homeostasis, which constrains neuronal firing rates within functional ranges. This regulation prevents pathological excitation-inhibition imbalances (Chen et al., 2022), such as hyperactivity or quiescence, thereby preserving the brain’s capacity to sustain diverse and flexible dynamics.

Biophysically detailed neuronal models are crucial for elucidating the mechanistic principles of neural function, enabling researchers to simulate and analyze how specific dynamics and synaptic interactions contribute to neuronal signaling and information processing. At a macroscopic level, many models have been used to explain the wide variety of global brain dynamics observed empirically (Cofré et al., 2020; Coronel-Oliveros et al., 2021; Herzog et al., 2023a; Luppi et al., 2023; Patow et al., 2024). Whole-brain models are pivotal in neuroscience as they enable the simulation of large-scale neural dynamics by integrating anatomical connectivity data with computational frameworks, providing insights into the mechanisms underlying brain functions. These models are instrumental in understanding how distributed brain networks interact to produce complex behaviors and cognitive processes. Models that incorporate plasticity mechanisms (in the form of local adaptation rules) can support a remarkably broad spectrum of neural dynamics using a single unifying mechanism (Hellyer et al., 2016; Rocha et al., 2018; Goldman et al., 2023; Stasinski et al., 2024; Rocha et al., 2024). For example, by strategically tuning these adaptive processes—such as homeostatic plasticity of excitatory/inhibitory connections or intrinsic neuronal excitability—they can reproduce both wake-like asynchronous firing and sleep-like slow-wave activity in a cohesive framework (Goldman et al., 2023). This way, complex shifts in global brain state can emerge naturally from local plastic changes, allowing networks to transition smoothly between different regimes while retaining stable yet flexible functional organization (Rocha et al., 2018; Stasinski et al., 2024). This built-in adaptability may be crucial not only to describe healthy conditions, like wakefulness and sleep, but also for investigating how pathological (Coronel‐Oliveros et al., 2024; Stefanovski et al., 2019) or pharmacologically altered states(Herzog et al., 2023a) arise when these mechanisms are disturbed.

While most such frameworks do not explicitly include homeostatic adaptation rules, they can still reproduce many different brain states by introducing targeted mechanisms for each phenomenon (Luppi et al., 2025). The Dynamic Mean Field (DMF) model (Wong & Wang, 2006; Deco, Ponce-Alvarez, et al., 2014a; Herzog, Mediano, Rosas, Luppi, Sanz Perl, et al., 2024) is a well-established example: through careful tuning of local excitability parameters and global coupling—often guided by empirical maps of neurotransmitter expression or other empirical brain heterogeneities (Deco et al., 2018; Herzog et al., 2023a; Luppi et al., 2025; Mindlin et al., 2024; Perl et al., 2023)—it has replicated the effects of pharmacological manipulations (Luppi et al., 2022; Herzog et al., 2023a) on brain-wide activity. Similarly, gradual changes in structural connectivity weights have been used to simulate aging(Gatica et al., 2022), by modeling the progressive decline of synaptic or axonal integrity. Schizophrenia-like states have been reproduced via increasing local self-excitation in specific subnetworks (Yang et al., 2016), while neurodegenerative disorders have been modeled through excitation/inhibition imbalances in regions undergoing atrophy (Moguilner et al., 2024) with specific hypo/hyperexcitability spatial patterns. Collectively, these parameter-based approaches highlight how a single model architecture can account for diverse observed states (Luppi et al., 2025), albeit typically without relying on a unified adaptation mechanism to maintain stable firing across conditions. However, this requires exhaustive parameter searches (Deco, Ponce-Alvarez, et al., 2014a) or restrictive heuristic solutions (Herzog, Mediano, Rosas, Luppi, Sanz Perl, et al., 2024) to set the local inhibition, also known as Feedback Inhibitory Control (FIC). Additionally, the solutions obtained are static, which can constrain free parameters and limit the model’s capacity to capture a broader range of observed brain dynamics.

As presented thus far, whole-brain models use individual mechanisms to control the dynamics that characterize a specific brain state. It is unclear how this approach can tackle conditions that are characterized by concurrent and distinct dynamical regimes. Dissociative states fall into this category, where asynchronous activity akin to wakefulness co-exists with slow waves typically present in sleep (Antelmi et al., 2016; Sodré et al., 2023). This paradoxical appearance of diverse dynamics bears resemblance to the notion of ‘chimeras’ from complexity science: when a subset in a system of identical parts, sustains disparate dynamics than the rest. In fact, chimeric states of brain dynamics are linked to cognitive functions (Bansal et al., 2019). Although the brain is not composed of identical parts since it features spatial gradients(Huntenburg et al., 2018) and use-dependent homeostatic regulation can alter dynamics in a regional way (Andrillon & Oudiette, 2023), we take inspiration from complexity science to call these dissociative states ‘chimeric-like’ and investigate how regional modulations of homeostatic processes underlie the emergence of concurrent distinct dynamical regimes.

Here, we embed a biologically grounded homeostatic inhibitory plasticity (Vogels et al., 2011) into the DMF framework, creating the Homeostatic DMF (HDMF). We then systematically chart the model parameter landscape to pinpoint an optimal solution that ensures homeostasis at the whole-brain level. Next, we test its adaptiveness and robustness when applying biologically plausible neuromodulation, revealing enhanced stability and predictability of the change on firing distributions. We then demonstrate that just by manipulating the global coupling the model exhibits sleep-like slow-wave activity (SWA), an unprecedented feature for this model. Furthermore, we achieve chimeric-like states by introducing spatially heterogeneous homeostasis. Finally, we show that the HDMF preserves the classic use of the DMF for resting-state fMRI fitting: it achieves comparable functional connectivity fits while improving the fit to functional connectivity dynamics. Importantly, this empirical fitting occurs in a wake-like regime, while sleep-like slow-wave activity remains accessible elsewhere in the HDMF parameter space. This demonstrates how a unified adaptation mechanism can reconcile diverse brain states in a single, cohesive framework.

## Results

We extended the Dynamic Mean Field model, a whole-brain approximation that Deco and colleagues introduced(Deco, Ponce-Alvarez, et al., 2014a), by incorporating a dynamic feedback inhibition control (FIC) mechanism with homeostatic plasticity, creating the HDMF model. The DMF model simulates neural activity by defining each region as consisting of interconnected excitatory (E) and inhibitory (I) neuron pools, with long-range interactions mediated solely between E pools via a structural connectivity matrix *C*_*ij*_ (here we used AAL cortical atlas Figure 1A, see Methods). A global coupling parameter *G* scales the influence of network interactions on each region. *G* is a free parameter used to fit empirical data by simulating blood-oxygenation-level–dependent (BOLD)-like signals from the excitatory firing rates time series *r*^*E*^ (Figure 1B). In its original version (Deco, Ponce-Alvarez, et al., 2014a), the DMF used a static feedback inhibition control (FIC) that required time-consuming optimizations to achieve stable firing rates, but a heuristic structure-dependent linear solution was later proposed bypassing this computational bottleneck (Herzog, Mediano, Rosas, Luppi, Sanz Perl, et al., 2024). This latter solution expresses the FIC as a function of the structural connectivity strength, *G* and another free parameter ***α***, which represents the global E/I balance (see Methods). However, both approaches are static in nature (i.e. implemented as constant scalar values), imposing strong limitations in terms of functional repertoire and firing rates stability especially in the high *G* (i.e. high coupled) regime. Our alternative dynamic solution for the FIC values regulates the excitatory firing rates around a target firing rate ρ based on a learning rate (LR) and a decay (*τ*_decay_) term (Figure 1A, B).

**Figure 1.**
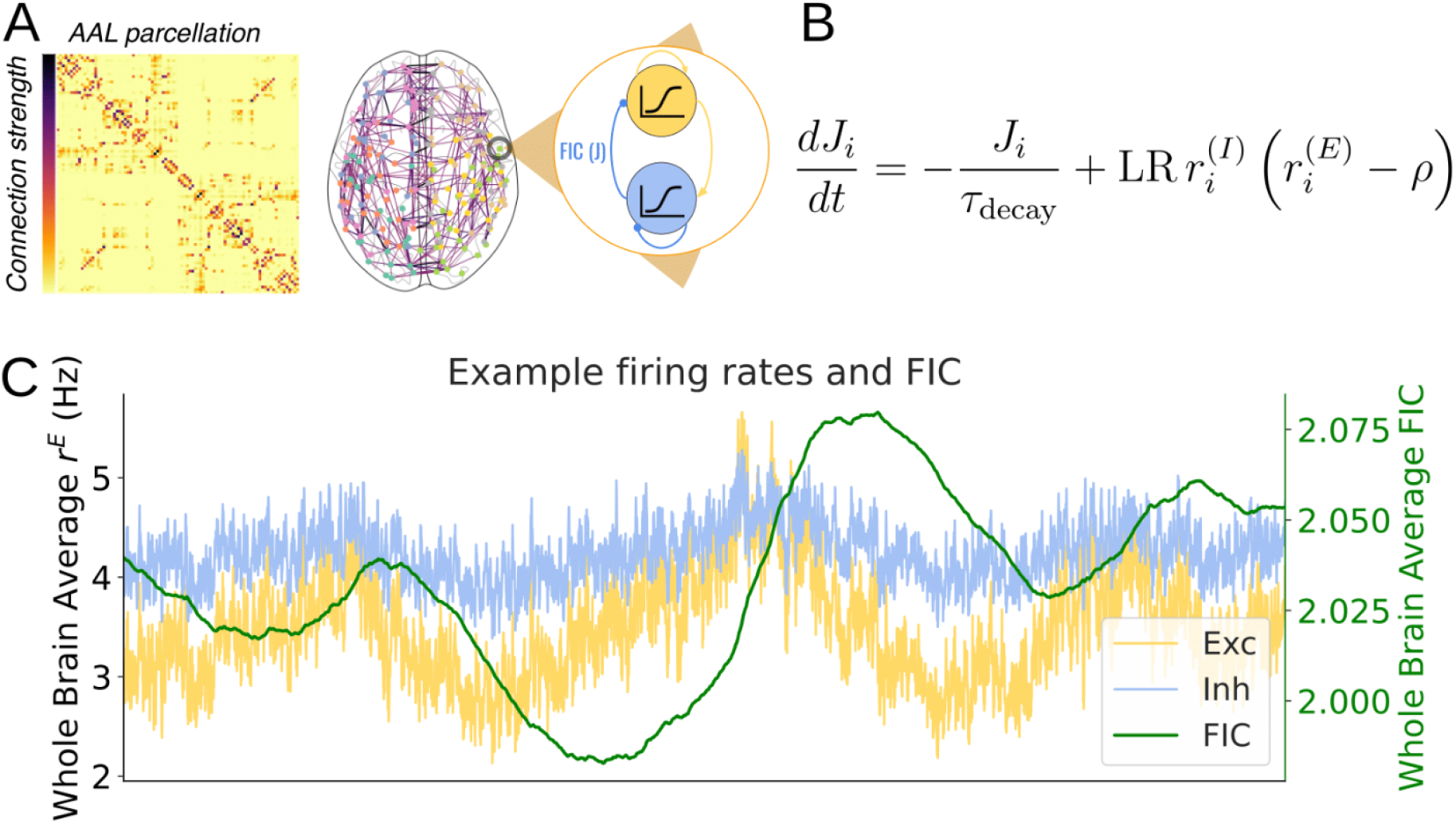
Overview of the Homeostatic Dynamic Mean Field (HDMF). **A)** Each brain region (circles) is defined by a parcellation and their connections by a structural connectivity matrix (edges) reconstructed from in vivo diffusion tractography. The dynamics of each region consists of reciprocally connected excitatory (E, yellow) and inhibitory (I, blue) neural populations, while inter-regional connections are only through the E population. **B)** The I->E connection, i.e the FIC variable *J*_*i*_ is controlled by a plasticity mechanism which dynamically adjusts the E firing rate (*r*^*E*^) to approach the target firing rate ρ depending on LR and *τ*_decay_. **C)** Example of time series of the simulated whole-brain average firing rates of an E (yellow) and I population (blue), and the respective average dynamic FIC (green) across all brain regions. Note that the FIC rapidly increases after sudden increases of the E firing rates, compensating for the excess firing rate, and thus reducing them. Note the delayed coupling between the E firing rates and the FIC.

### Homeostatic dynamic mean field model (HDMF)

Here, we aimed to identify the regime in which an inhibitory plasticity rule enables effective homeostatic control within the Dynamic Mean Field model, which we refer to as the HDMF. To this end, we computed the homeostatic fit, measured as the percentage mismatch between the average excitatory firing rate and the target firing rate (ρ = 3.44 Hz (Deco, Ponce-Alvarez, et al., 2014a)), as a function of the learning rate (LR = [10^0^, 10^3^] ms^-1^) and the decay (*τ*_decay_= [10^2^, 10^5^] ms) (Figure 2A). To demonstrate the enhanced stability, the average mismatch was computed across a wide range of G values, from 0 to 8.5. For each fixed value of *τ*decay, we identified the LR that minimized this mismatch; these minima are shown as green dots in Figure 2A. Thus, the green dots mark the LR–*τ*decay combinations for which the plasticity rule best controls the firing rate around the target. Since the results are plotted in log-log scale, the alignment of these minima confirmed the power-law relationship between the learning rate and decay that optimizes homeostasis, obtained analytically (see Methods: Analytical derivation of the LR-τdecay power-law relationship). The estimated exponent closely approaches the theoretical values obtained in an analytical derivation. Further, we found that the region of parameters for optimal homeostasis also corresponds to minimal variability in firing rates across the range of G, indicating enhanced stability regardless of the G value (Figure 2B) and target firing rates (Supplementary Figure 1). Note that without the decay term the average firing rate never converged to the target firing rate (Supplementary Figure 2). Then, in the optimal regime, the homeostatic plasticity rule can be expressed only in terms of the learning rate and ρ, as *τ*_decay_ can be derived from the power-law fit. In biophysical terms, this relationship implies that the faster the synaptic weight is learnt (high LR), the faster it needs to be forgotten (fast decay; small *τ*_decay_) in order to adapt to the ongoing fluctuations of network inputs. From here on, whenever we use the HDMF, it means that the *τ*_decay_ is derived for a fixed LR and ρ using the power-law relation from the inset in Figure 2A.

**Figure 2.**
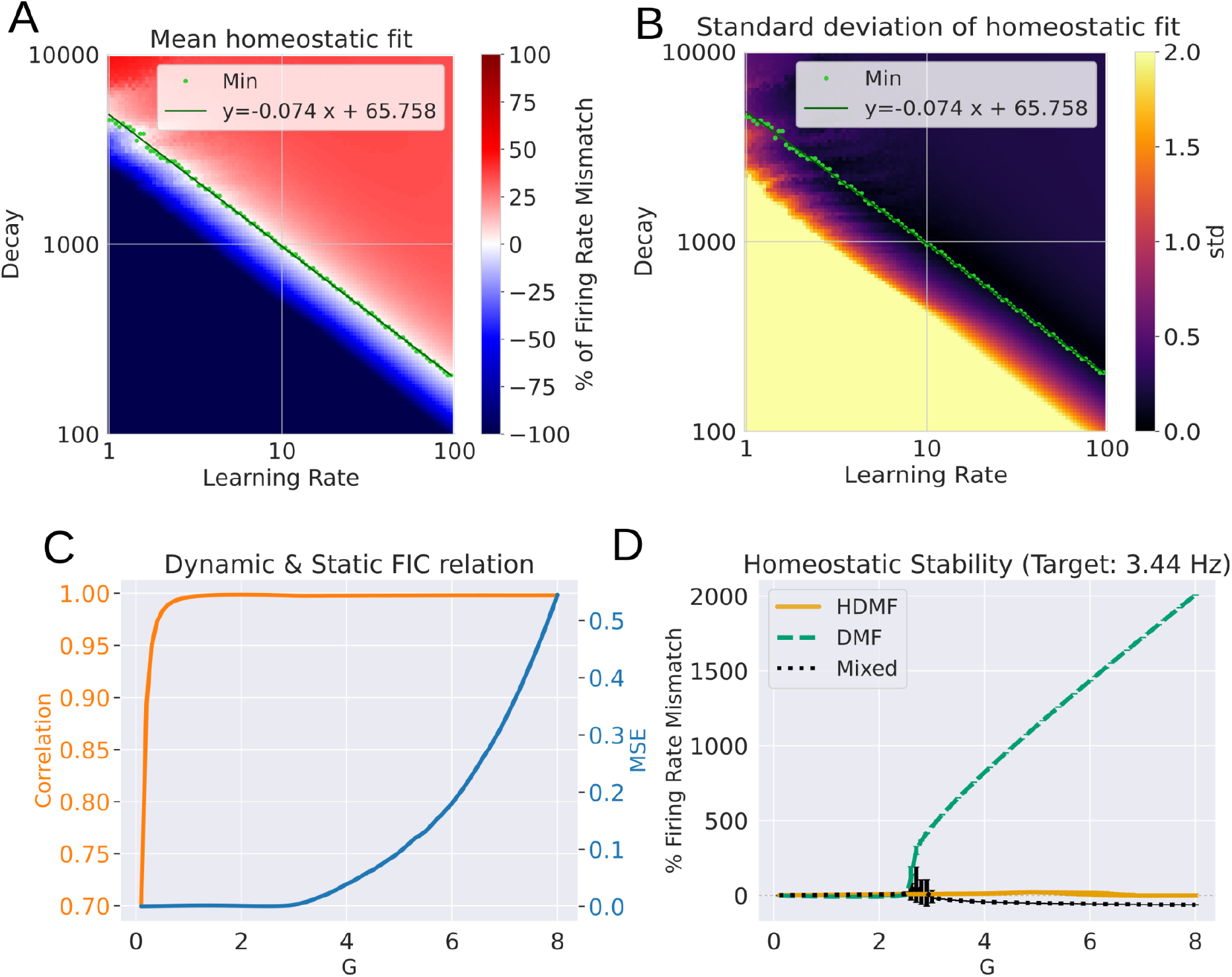
The local inhibitory plasticity controls whole-brain homeostasis. **A)** Average homeostatic fit (the percentage of mismatch between the average E firing rate and ρ) as a function of the learning rate and the decay for ρ= 3.44 Hz in log-log scale. Averages were taken across a range of G values from 0 to 8.5. Green dots show, for each fixed *τ*decay value, the LR that minimizes the mismatch between the mean firing rate and ρ. The darker lines show the linear fit to these minima in log-log space, corresponding to the power-law relationship between LR and *τ*decay used to define the homeostatic regime. Note that since the plot is in log-log scale, this relationship is a power-law. The coefficients of the linear equation inset in the panel are used to implement the HDMF at the desired target firing rate. **B)** Same as in **A**, but showing the standard deviation in the same range of G values. This shows that the region of minimal mismatch is also the region with more stability across G values. **C)** Correlation and the mean squared error (MSE) between the static FIC obtained by the linear solution (see Methods) and the one obtained after averaging the dynamic FIC over time, as a function of G. Note also that close to G=2.5 the MSE diverges, meaning that beyond this point the linear solution for the FIC is no longer the time-average of the dynamic FIC. Despite their high correlation, a static FIC is not able to achieve stability, demonstrating the relevant role for the dynamic FIC in achieving firing rate stability. **D)** Percentage of firing rate mismatch as a function of G for the DMF (Static, green dashed line), for the HDMF (Dynamic, yellow line) and for a DMF where the FIC values corresponded to the average of the dynamic FIC (Mixed, black dots). Note that close to G=2.5 the DMF diverges into a high-firing rate regime, while the HDMF maintains the activity close to the target. The mixed model could not sustain stability near this divergence point. For higher G values, the mixed model stabilized but below the target rate, indicating over inhibition. For panels **C** and **D**, the DMF used ***α***=0.75 while the HDMF used LR=3.5. Both were simulated for 50000 time points using the same set of 30 different initial conditions. Lines and shadowed areas represent the mean and standard deviation across initial conditions, respectively.

To investigate the nature of the FIC values obtained through plasticity, we compared the average firing rates as a function of G for three different models. First, the original DMF model that uses a static FIC for each region, derived from a linear solution (Herzog, Mediano, Rosas, Luppi, Sanz Perl, et al., 2024) or using pseudo gradient-descent algorithms (Deco, Ponce-Alvarez, et al., 2014a). Second, the HDMF that dynamically changes FIC values to sustain a constant average firing rate. Notably, one might expect that, when averaged over time, the time-varying FIC of the HDMF could converge toward FIC values similar to those obtained through static methods. Thus we also created a third model, the “mixed” DMF, that uses the FIC time-average values found from the HDMF but does not dynamically adjust these values. For the same range of G values, we computed the correlation and mean squared error (MSE) between the static FIC values obtained from the linear solution and the time-averaged dynamic FIC values (Figure 2C). At very low G values—corresponding to weak network input—the FIC dynamics were predominantly noise-driven, reducing the correlation between the static and dynamic FIC values in this regime. For the remaining range we found that while the correlation remained maximal, the MSE markedly increased near G=2.5, indicating a difference between static and dynamic FIC values in this regime. Consistently with previous results (Deco, Ponce-Alvarez, et al., 2014a; Herzog, Mediano, Rosas, Luppi, Sanz-Perl, et al., 2024), the DMF diverged into a high-firing rate regime near G=2.5 whereas the HDMF maintained firing rates close to the target across the entire range of G values (Figure 2D). Additionally, despite using FIC values derived from the HDMF, the mixed DMF model failed to achieve stability near this critical point and remained well off the target rate -40% mismatch-for higher G values. This highlights the necessity of dynamic FIC to compensate for rapid fluctuations in firing rates.

Overall, these results demonstrated that the dynamics of the FIC, beyond the simple average, were necessary for achieving stability in a wide range of G. In fact, in the low-to-mid coupling regime (0.5<G<2.5) the heuristic linear solution for the static FIC is a particular case of the dynamic FIC, suggesting that the dynamics of the latter could be represented by its steady-state value within this regime. Beyond this regime, this relationship does not hold anymore.

### Enhanced stability under neuromodulation

We investigated the stability of the HDMF model under neuromodulation, using the receptor map of the 5HT-2A receptor obtained from positron emission tomography (PET) images (Hansen et al., 2022), which is widely expressed in the cortex. Neuromodulation was simulated via response gain modulation (Deco et al., 2018), where an increase in the neuromodulatory gain elevates the slope of the F-I curve in both E and I pools (Herzog et al., 2023a), resulting in higher firing rates with the same input current (Figure 3A). As each region will have a specific value of receptor expression, this mechanism will impact regions differently. A major obstacle of this approach in the DMF is that it largely impacts the firing rates, thus losing biological plausibility for high neuromodulatory gains. This is critical because the resting-state DMF formulation assumes that each cortical region operates near a low-firing spontaneous regime, maintained around the ∼3 Hz working point by feedback inhibition control. Functional connectivity is then generated by noisy fluctuations around this low-activity fixed point (Deco, Ponce-Alvarez, et al., 2014a). Therefore, neuromodulatory perturbations that drive firing rates far above this regime push the model away from the mechanistic operating point for which the DMF was designed.

**Figure 3.**
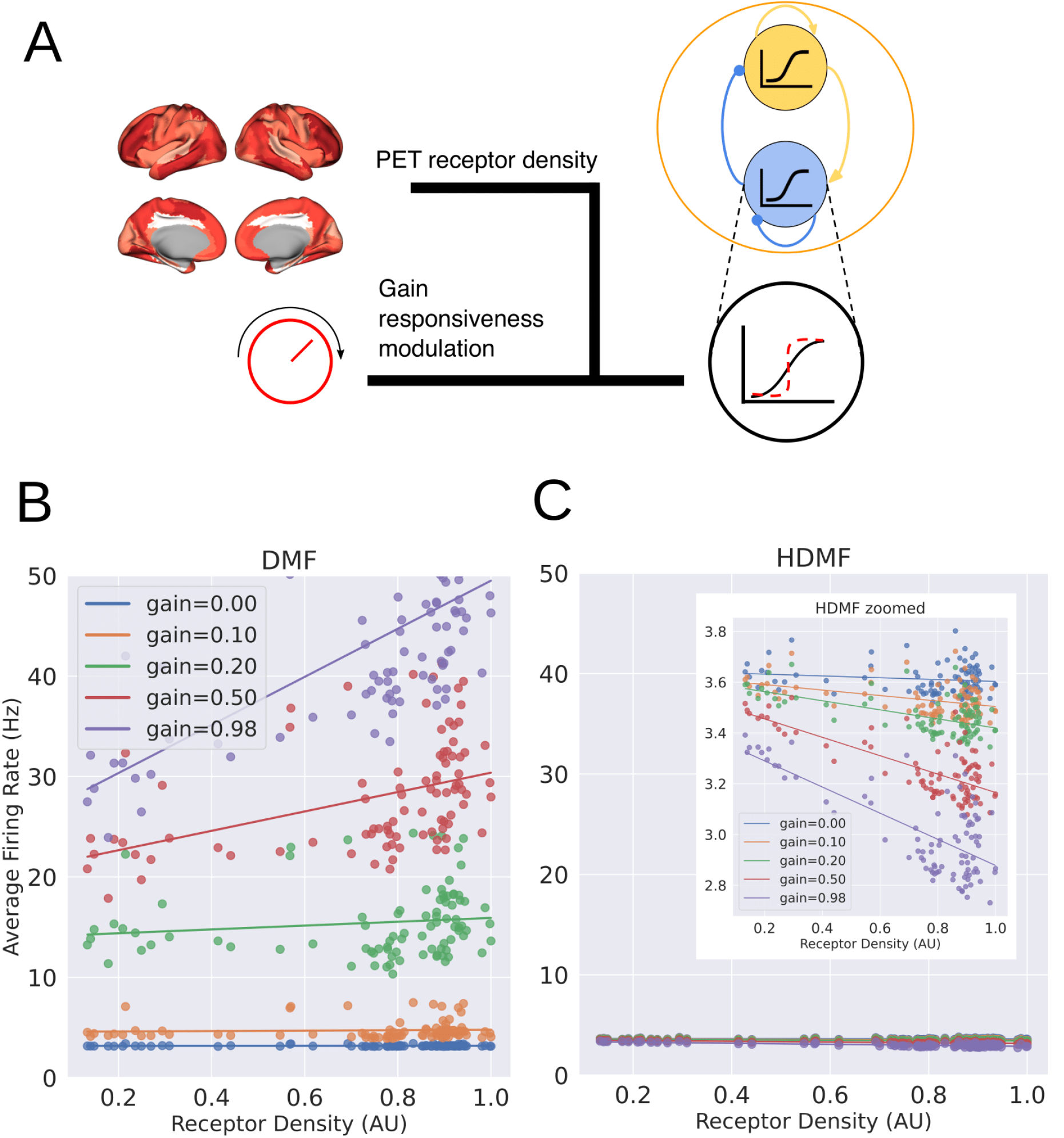
Homeostasis stabilizes the effect of increasing neuromodulation. **A)** Schematic representation of neuromodulation through response gain modulation in the simulation. As the neuromodulatory gain increases, the slope of the F-I curve increases, evoking higher firing rates with the same input current. **B)** Average firing rate vs the receptor density for the DMF. Each dot represents a brain region. Colors denote 5 different neuromodulatory gains. Increasing neuromodulatory gain drives the DMF away from the low-firing spontaneous regime, producing unrealistically high firing rates across regions. At high gains, the receptor-density dependence is therefore dominated by runaway excitation rather than by a biologically interpretable regional modulation. **C)** Same as **B**, but for the HDMF. In contrast, the HDMF preserves firing rates near the low-activity working point targeted by feedback inhibition control, despite increasing neuromodulatory gain. This indicates that regional receptor-density effects can be expressed without pushing the model outside the biologically plausible spontaneous-activity regime. Both models were simulated for 100 different initial conditions. DMF used ***α***=0.75 and G=2.11; HDMF used LR=3.5 and G=2.55.

Here, we compared the firing rate stability of the DMF model with the HDMF model in response to increasing neuromodulatory gains. In the standard model, firing rates increased with gain, but at high gain values, the positive correlation with receptor density was highly variable, showing a wide spread around the regression line (Figure 3B). In contrast, the homeostatic model maintained firing rates closer to the target (3.44 Hz), even with large neuromodulatory gains. Furthermore, the changes in firing rates in the HDMF model exhibited a tight negative dependence on receptor density, demonstrating a more predictable and controlled response (Figure 3C). Notably, this negative correlation is the result of the homeostatic process: as more gain implies amplified responses for the same input currents, the departure from the target firing rate will also amplify the dynamic feedback inhibition control response, driving the excitatory firing rate down on average in consequence. However, this offset was minor, in comparison to the explosive effect observed with the DMF, and still provides an enhanced range for simulating neuromodulation with biologically plausible firing rates. This stability was not limited to a particular gain value or neurotransmitter distribution (Supplementary Figure 3), underscoring that the HDMF supports a broader and more robust neuromodulatory regime than the DMF. By contrast, a mixed model (as in the previous section) used as a control yielded inconsistent outcomes (Supplementary Figure 3), supporting again that the dynamics of the FIC provide a stability that its steady-state value alone cannot achieve. We further found that this stability was preserved when neuromodulatory gain varied continuously over time: under dynamic gain, the HDMF maintained firing rates close to the target regime despite transient gain-driven excursions, whereas the DMF entered a sustained overexcited state (Supplementary Figure 4)

Finally, we tested whether this enhanced stability under neuromodulation preserved the model’s ability to fit empirical pharmacological data. To this end, we reproduced the analysis of Herzog and colleagues (Herzog et al., 2023b) using fMRI data from participants under LSD, a 5-HT2A receptor agonist (Supplementary Figure 5). At the optimal fitting points, the HDMF maintained firing rates within a narrower and more biologically plausible range than the DMF, while also providing better fits to the empirical data. Thus, the homeostatic mechanism improves stability under receptor-informed neuromodulation without reducing empirical fitting performance, and in this case improves it.

### A potential mechanism for whole-brain slow wave generation

Recent advances in whole-brain modeling aim to reproduce different brain states through single parameter manipulations thus providing insightful and interpretable mechanisms about the transitions of brain dynamics (Luppi et al., 2025). Slow waves, whose underlying bistable dynamics alternates periods of neural activity and silence (Sanchez-Vives, 2020), are central for diverse biological and cognitive functions (Sheybani et al., 2023; Wilckens et al., 2018) and are present in states such as deep sleep, general anesthesia and disorders of consciousness (Sanchez-Vives, 2020). Previous attempts to generate slow wave activity (SWA) in whole-brain models have employed different mechanisms (Goldman et al., 2023; Marsh et al., 2024; Sacha et al., 2025) but it has never been done in the DMF, although this model has been widely used to study different states of consciousness.

Here, we aimed to demonstrate that the homeostatic model can generate slow-wave activity through modulation of the learning rate LR parameter. Given a high global coupling G, as LR increased, the whole-brain average of the excitatory firing rates transitioned from fully regular SWA to irregular activity (Figure 4A). This transition was further captured by the emergence of pronounced peaks in the power spectra at frequencies consistent with empirical delta slow waves (1-2 Hz), confirming the presence of periodic activity, opposed to the flat power across frequencies characteristic of irregular regimes (Figure 4B).

**Figure 4.**
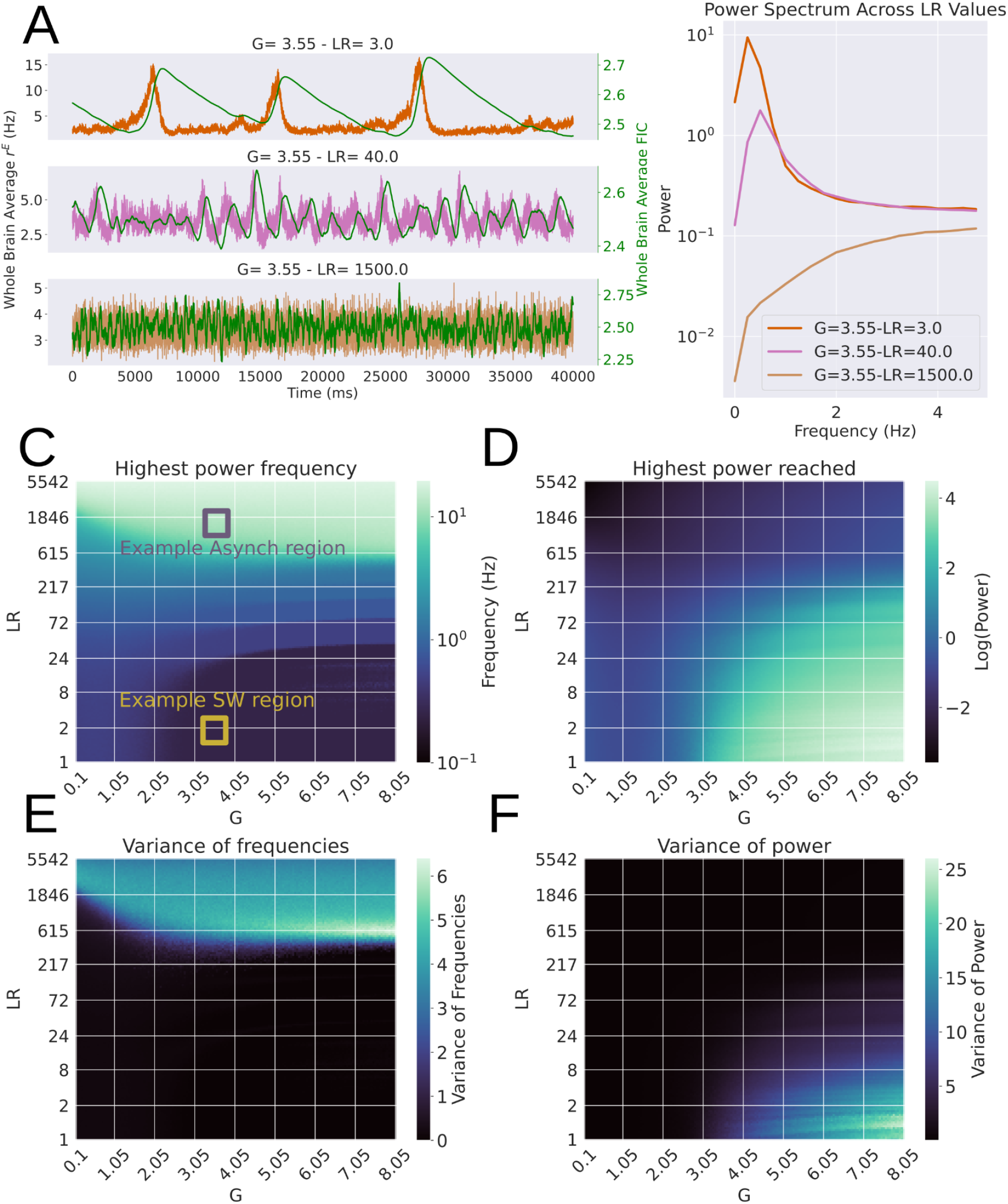
Slow waves emerge in the HDMF in the highly coupled regime. **A)** Whole-brain average excitatory firing rate for three different learning rate values (LR), with fixed high global coupling (G=3.55). **B)** Power spectra of the time series shown in A. Low-frequency power is prominent in the slow-wave regime and decreases as LR increases, consistent with the transition toward irregular activity. **C)** Average maximal frequency across regions as a function of LR and G. Colored squares show areas where points are taken for Fig 5A. **D)** Average power of the maximal frequencies shown in **C**. Values in C and D are averages out of 8 different initial conditions. **E** and **F)** Standard deviations for (C) and (D), respectively, across initial conditions. Note the increased variability near the transition between the slow-wave and irregular activity regimes.

**Figure 5.**
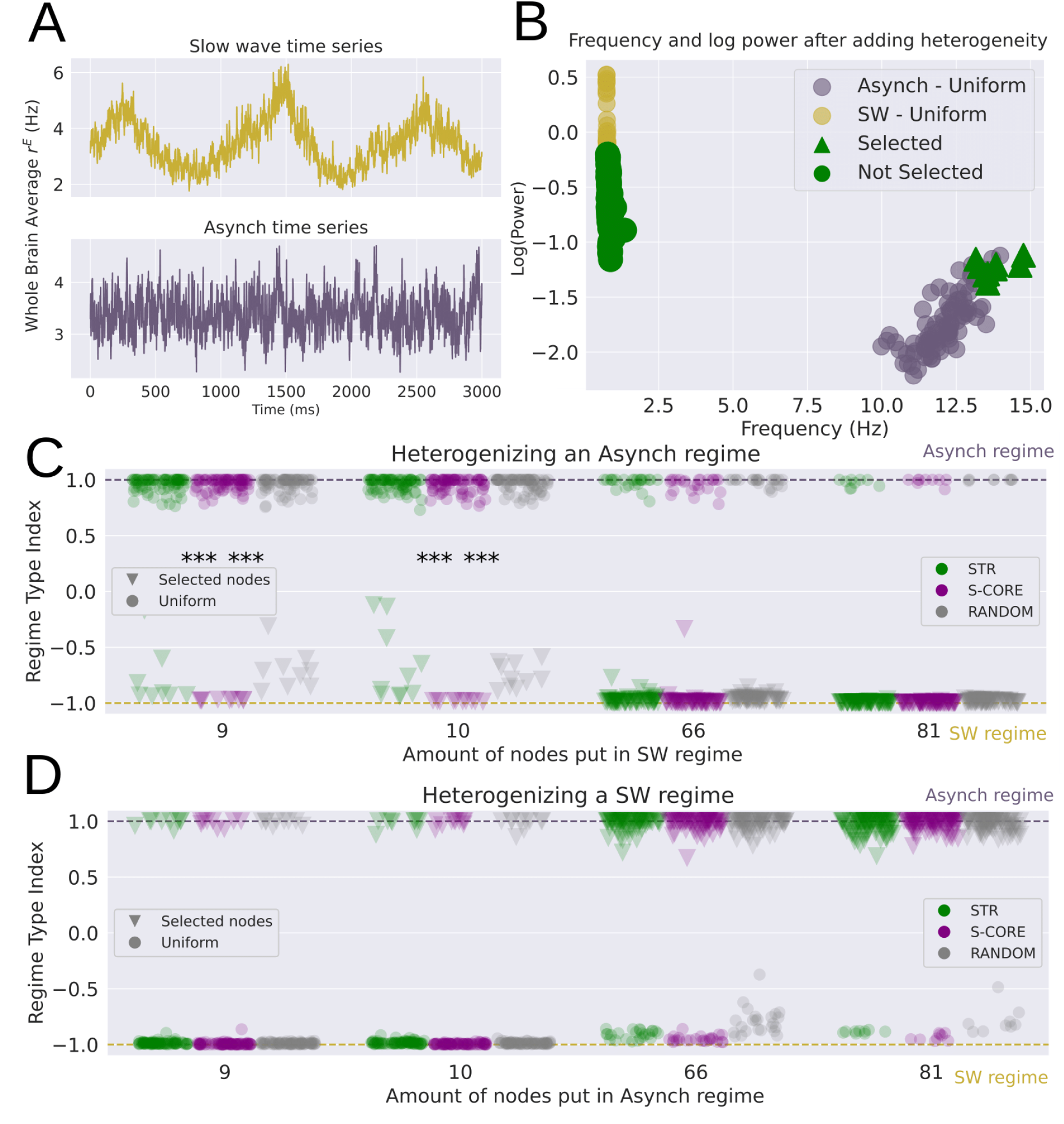
Regional heterogeneity in the learning rate (LR) gives rise to chimeric-like dynamics. **A)** Uniform baseline simulations at high global coupling (G = 3.55). A low learning rate (LR = 100) generates slow-wave activity, whereas a high learning rate (LR = 1,500) generates asynchronous dynamics. **B)** Example heterogeneous simulation starting from the slow-wave regime. Nine STR-selected regions were assigned high LR (green triangles), while the remaining regions retained low LR (green circles). Dots show the uniform slow-wave baseline (yellow) and the uniform asynchronous baseline (grey). Selected high-LR regions shifted toward the asynchronous frequency–power regime, while non-selected low-LR regions remained closer to the slow-wave regime but showed reduced power. **C)** Asynchronous-to-slow-wave manipulation. Starting from a uniform high-LR asynchronous regime, selected regions were assigned low LR. The Regime Type Index (RTI) quantifies each region’s proximity to the two uniform baselines, with RTI = −1 indicating the slow-wave regime and RTI = 1 indicating the asynchronous regime. For small selected groups, regions reached slow-wave-like RTI values mainly when they formed an S-CORE subnetwork; STR and random selections produced only partial shifts. **D)** Slow-wave-to-asynchronous manipulation. Starting from a uniform low-LR slow-wave regime, selected regions were assigned high LR. In this direction, selected regions reached asynchronous-like RTI values across group sizes and selection criteria. Group sizes were chosen to match the subnetwork sizes obtained with the S-CORE algorithm. *** indicates Cohen’s d > 0.8

We then systematically explored the power spectra across a wide grid of LR and G values, identifying for each parameter combination the average dominating frequency across all regions and their corresponding power. This analysis revealed a broad landscape of delta band frequency (Figure 4C) with high power (Figure 4D). As we increase the LR, frequency increases and power decreases, as a consequence of faster learning dynamics. This reaches a critical point where variability peaks and the system enters a general high frequency with lower power dominated state (Figure 4E-F). These findings reveal a potential whole-brain mechanism of slow wave generation within the HDMF model. The homeostatic plasticity is at the core of this mechanism: for the high-coupled regime, the excitatory and synchronized inputs from the network abruptly increase a given region’s firing rate, with the consequence of an abrupt rise in the FIC. As a consequence, the firing rates decrease rapidly as well and remain down until the FIC decays enough to trigger another active state. When LR is too high, the compensation is much faster, generating more noise driven irregular dynamics with less dominance of slow frequencies. The capacity to induce SWA through a biophysically grounded homeostatic rule highlights the HDMF model’s versatility and interpretability in capturing diverse brain states.

### Heterogeneous plasticity enables different and coexistent dynamics

Until now we explored the new dynamical repertoire of the DMF model when incorporating a homeostatic plasticity rule, which facilitates the emergence of SWA. However, this plasticity was uniformly applied across all regions, leading to fully synchronized dynamics. In contrast, biological plasticity is inherently heterogeneous, varying across the brain (Appelbaum et al., 2023; Huntenburg et al., 2018) –giving rise to distinct roles and computational capacities. Since the HDMF can self-regulate, we asked whether it could simultaneously sustain different dynamical regimes via heterogeneous plasticity. Motivated by the aforementioned chimeric-like states, we studied what happened to a model set in a system-wide asynchronous regime, when we change adaptiveness through the learning rate of some regions to induce in them a regional slow-wave activity regime, and, conversely, starting from system-wide SWA, shifting regions toward an asynchronous regime.

We created baseline simulations at fixed high coupling (G=3.55): a uniform low LR (LR=100; SWA regime) state and a uniform high LR (LR=1,500; asynchronous regime) (Figure 5A). To select which regions to change their LR value with respect to the starting global uniform LR value we used three criteria based on anatomical connectivity, namely node strength (STR) and strength-core (S-CORE), and random regions (RANDOM) (Castro et al., 2020; Rubinov et al., 2009). Node strength is a local graph theoretical measure that corresponds to the sum of all the connection weights for a given region. S-core is a mesoscopic measure aiming to detect the maximal subgraph in which every node has strength -at least-*s*. When using node-strength we choose the *k* regions that have the highest average connectivity. For strength-core we select the *k* regions that have the same or higher connectivity that are also connected between each other.

To characterize each region’s local dynamics, we computed its peak frequency and corresponding power. The uniform asynchronous regime showed higher peak frequencies and lower power, whereas the uniform slow-wave regime showed lower peak frequencies and higher power. Figure 5B illustrates one heterogeneous example in which selected regions were assigned high LR within an otherwise low-LR slow-wave background. The selected regions shifted toward the asynchronous regime, while the remaining low-LR regions stayed closer to the slow-wave regime but with reduced power, reflecting the interaction between both dynamical subgroups. To quantify this shift we define the Regime Type Index (RTI) which measures proximity to each baseline frequency (see Methods for details). For an RTI of 1, the region’s peak frequency matches the asynchronous regime, while for an RTI of -1, it matches the SWA regime. Inspired by chimera states studied in complex systems theory, we call the co-existence of regions with both high and low RTI values a chimeric-like state.

We repeated this procedure for increasing numbers of selected regions and for both directions of manipulation (Figure 5C–D). Triangles represent regions selected to have a different LR regime. When starting from the asynchronous regime and switching selected regions to low LR, the emergence of slow-wave-like dynamics depended on the topology of the selected group. STR-selected and randomly selected regions shifted only partially toward the slow-wave regime, whereas S-CORE-selected regions reached the lowest RTI values, consistent with a fully slow-wave-like regime. This difference was strongest for groups of 9 and 10 regions. In contrast, when starting from the slow-wave regime and switching selected regions to high LR, selected regions reached asynchronous-like RTI values across group sizes and selection criteria. Thus, inducing local slow-wave activity within an asynchronous background was topology-dependent, whereas inducing local asynchronous activity within a slow-wave background was comparatively robust.

### HDMF fit to resting state fMRI data

After showing the novel features of the HDMF in this section we confirm that these modifications do not affect classic use of the DMF to reproduce slow fluctuating dynamics. We aimed to compare the DMF model (with the heuristic linear solution for the FIC) with the HDMF model in terms of their fit to empirical resting state fMRI data. To this end, we used resting state fMRI data from 99 unrelated HCP-YA subjects and ran the complete pipeline from end to end, calibrating the learning rate-τ_decay_ relationship for this specific parcellation and connectome (Supplementary Figure 6). To fit empirical data, we computed two observables: i) the functional connectivity (FC, Figure 6E) matrices, and ii) the functional connectivity dynamic (FCD) histograms (Figure 6F). These observables have been typically used to fit whole-brain models to fMRI data (Herzog et al., 2023a; Herzog, Mediano, Rosas, Luppi, Sanz Perl, et al., 2024; Luppi et al., 2025), as they can be reliably estimated from the usual sample sizes of fMRI recordings. We explored parameter space using a grid of 31 values of the global coupling G (0.0 to 3.0 in steps of 0.1) and 11 values of the excitation-inhibition balance parameter α (0.0 to 1.0 in steps of 0.1) for DMF, and 51 values of G (0.0 to 5.0 in steps of 0.1) and 41 logarithmically spaced values of the learning rate (LR, ranging from 0.1 to 1000.0) for HDMF. For each parameter combination, we ran 20 simulations with identical seeds for both models and computed both observables. Goodness of fit was quantified using the correlation between the population-averaged empirical FC and the simulated FC, and the Kolmogorov-Smirnov (KS) distance between the pooled empirical FCD histogram and the simulated FCD histogram (Deco et al., 2018). As shown in Supplementary Figure 7, the parameter regions yielding optimal fits for FC and FCD partially overlapped for both models. To identify parameter configurations that fit both observables, we selected the top-performing parameter sets for FC and FCD separately and retained only those belonging to the intersection of the two sets (Figure 6A-B). We first pooled the seeds to obtain the population-averaged FC and the pooled FCD histogram. We then identified the first non-empty overlap region, which emerged at 10%. Within this region, the optimal DMF parameters were G = 1.0 and α = 0.7, yielding an FC correlation of r = 0.577 and an FCD KS distance of 0.399. The corresponding HDMF optimum was G = 1.1 and LR = 0.8, yielding an FC correlation of r = 0.563 and an FCD KS distance of 0.290. We used these parameters for running 100 simulations with the same seeds and robustly compared the models’ performances.

**Figure 6.**
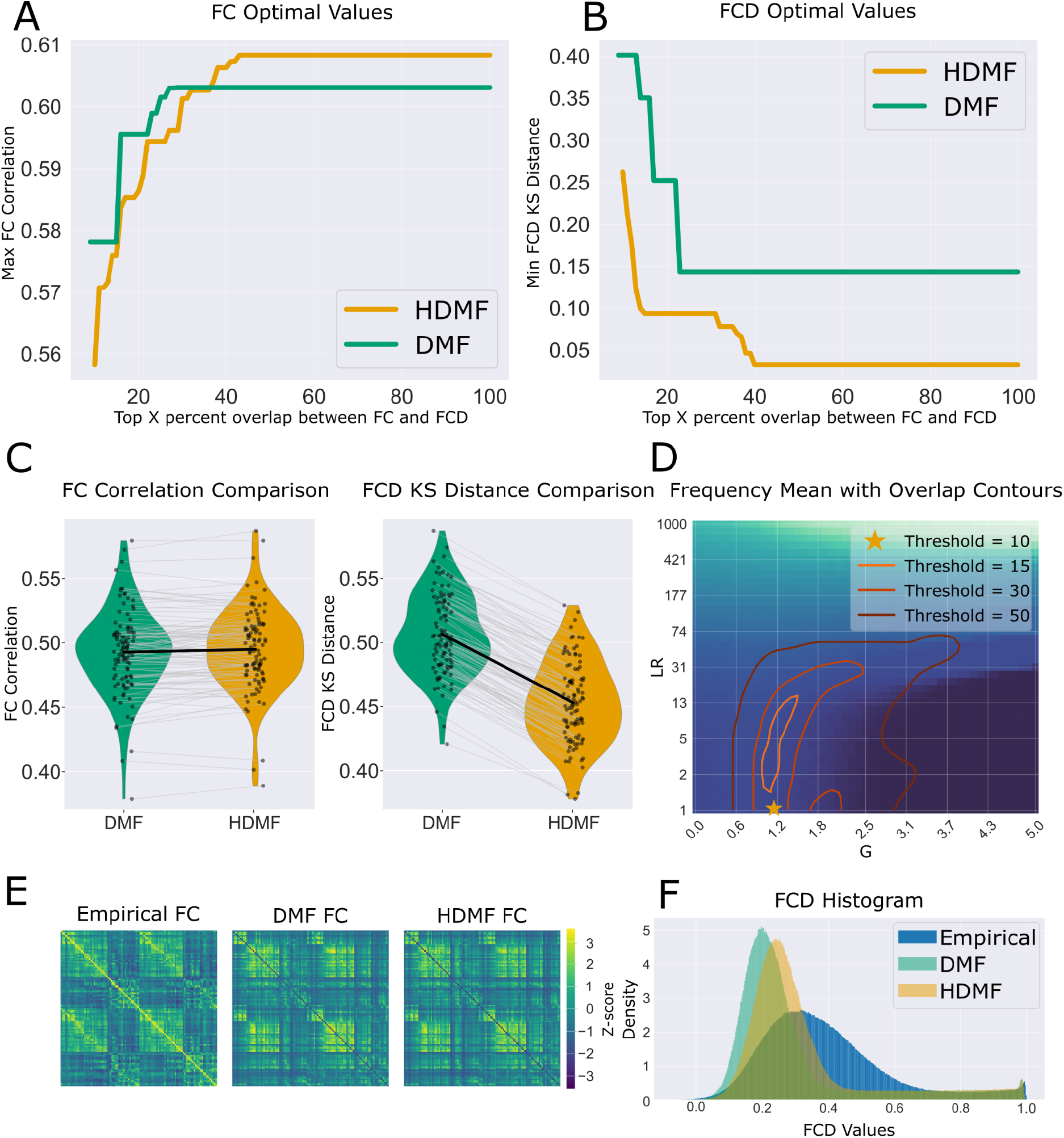
Empirical resting-state fMRI fit of the DMF and HDMF models. **A)** Maximum FC correlation obtained within the overlap between the top-performing FC and FCD parameter sets, shown as a function of the top X% tolerance threshold. Curves are shown for the static-FIC DMF and the HDMF. **B)** Minimum FCD Kolmogorov–Smirnov (KS) distance obtained within the same overlap regions. For both models, the first non-empty FC/FCD overlap emerged at the 10% threshold, which was used to select the joint-fit parameters. **C)** Paired model comparison across 100 simulations run with identical seeds at the selected parameters. Left: FC correlation with the empirical FC. Right: FCD KS distance. Grey lines connect paired simulations with the same seed, and black lines indicate the mean. The HDMF preserves a similar FC fit while reducing the FCD KS distance relative to the DMF. **D)** HDMF parameter-space map showing the mean dominant frequency as a function of global coupling G and learning rate LR. Contours indicate FC/FCD overlap regions obtained with increasingly permissive thresholds. The star marks the selected joint-fit point, which lies outside the slow-wave activity regime. **E)** Empirical FC matrix and simulated FC matrices for the DMF and HDMF at the selected joint-fit parameters. **F)** Pooled FCD value distributions for empirical data, DMF, and HDMF.

Across the 100 paired seeds of the 600 s run, FC correlation was very similar between models, with a marginally higher value for the HDMF than the DMF (HDMF: r = 0.495 ± 0.032, 95% CI [0.489, 0.501]; DMF: r = 0.493 ± 0.033, 95% CI [0.486, 0.499]), with a negligible paired effect (Cohen’s d = −0.21, computed on DMF − HDMF paired differences). FCD KS distance was lower for the HDMF than the DMF (HDMF: KS = 0.306 ± 0.064, 95% CI [0.293, 0.319]; DMF: KS = 0.413 ± 0.062, 95% CI [0.401, 0.425]), with a large paired effect (Cohen’s d = 4.66, computed on DMF − HDMF paired differences). This paired comparison is shown in Figure 6C, with the corresponding average FC matrices and pooled FCD distributions shown in Figure 6E–F. Although the FC fit is very similar, the HDMF shows a clear improvement in FCD, suggesting that the plasticity extension mainly benefits the temporal structure captured by FCD while preserving the static FC fit. Figure 6D demonstrates how this optimal point in the overlap of FC and FCD lies outside of the Slow Wave Activity regime, and how overlap region increases as we select more top-performing parameter sets.

To go beyond homogeneous firing rates and, consequently, homogeneous homeostatic constraints, we allowed the target firing rate to vary across brain regions in the HDMF. Starting from the previously identified optimal G and LR that maximized FC correlation within the first non-empty overlap region, we used Bayesian optimization to fit a heterogeneous target-rate vector across regions (Supplementary Figure 8). This yielded a stronger FC fit than the homogeneous-rho model, increasing the population-averaged correlation from r = 0.562 to r = 0.642. When using exactly the same 100 seeds, the FC correlation was higher for heterogeneous targets than for homogeneous ones (Pearson r = 0.541 ± 0.024, 95% CI [0.537, 0.546] and r = 0.495 ± 0.032, 95% CI [0.489, 0.501], respectively), with a large paired effect (Cohen’s d = 1.21). These results indicate that allowing heterogeneous homeostatic targets improves data fit while moving the model closer to biologically plausible region-specific homeostatic constraints.

## Discussion

### Overview of Main Contributions

One of the brain’s astonishing features is the ability to switch between radically different cognitive states while maintaining baseline functions. This robustness is partly grounded in its capacity to adapt to external and internal fluctuations. In this study, we introduced a homeostatic plasticity rule into the DMF model to enhance its biological plausibility and prove how this can support a broader repertoire of dynamics. This self-tuned regulation led to a robust maintenance of firing rates in different scenarios such as high coupling or following changes after neuromodulatory drive. The plasticity rule also allows for explicit selection of target firing-rate, which was only accessible in the DMF by indirect tuning of the Feedback Inhibition Control parameter. In addition, the HDMF generated slow waves—a phenomenon not previously captured by the DMF—and supported the coexistence of multiple dynamical states within the same system, thereby enabling the modeling of dissociative conditions such as parasomnias (Antelmi et al., 2016).

### Relevance of Homeostatic Plasticity for Whole-Brain Modeling

We first tuned the plasticity rule to self-regulate (i.e., reach homeostasis with respect to the model’s target firing rates). We achieved this by finding an analytical power-law relationship between the two control parameters, that depends both on the target firing rate and the specific connectivity matrix. We obtained these parameters, and confirmed that simulated firing rates reached the target rate with a neglectable deviance. The scaling reflects a generic constraint of homeostatic plasticity rules with linear decay rather than a property of HDMF specifically; its biological relevance lies in linking the timescale of synaptic forgetting to the learning rate required to maintain a target firing rate, with coefficients that depend on the underlying network and target activity. The central premise of homeostatic plasticity is to regulate local or global excitability, preventing runaway dynamics or quiescence (Abeysuriya et al., 2018; Soldado-Magraner et al., 2022). Integrating such a mechanism into the DMF –a well established biophysical whole-brain model– offers a biophysically grounded means to self-tune neuronal activity, thereby reducing sensitivity to precise parameter configurations. Previous solutions for calculating FIC include linear heuristics (Herzog, Mediano, Rosas, Luppi, Sanz Perl, et al., 2024) and pseudo‐gradient optimization(Deco, Ponce-Alvarez, et al., 2014a). The former relies on a relationship between structural connectivity and global coupling—providing mechanistic insight into E/I balance that is useful for spontaneous activity but falls short under extrinsic perturbations such as neuromodulation or external stimuli. In contrast, region‐specific optimization overcomes this limitation by using short simulations to adjust FIC values until stability is reached, thereby adapting weights to any modeling condition. However, this approach incurs high computational cost and compromises biological realism, as the brain can’t compute its synaptic weights in advance. Within the typical range of G values used in previous studies (0-2.5), the average FIC values produced by our rule match those obtained with previous methods. This highly effective self-regulation may not be present in pathological states. This opens an intriguing avenue for future exploration—one that first requires us to lay the groundwork. Accordingly, we have integrated homeostatic regulation into the fastDMF framework (Herzog, Mediano, Rosas, Luppi, Sanz Perl, et al., 2024), in keeping with its mission to simplify model development, ensure accessibility and move toward canonical models whose empirical validation can be more fully explored.

By adding the homeostatic plasticity rule, the HDMF generated slow waves. In this regime the system exhibits bistable dynamics, rapidly alternating between states of synchronous activity (UP) and synchronous silence (DOWN). When global coupling was high, slow adaptiveness gave rise to slow waves, whereas fast adaptiveness generated noisy activity. This coupling-adaptation mechanism is compatible with the low-level mechanisms driven by activity-dependent potassium currents (Compte et al., 2003). Additionally, other modelling studies propose that this mechanism generates a bistable state between two attractors (Holcman & Tsodyks, 2006). In vivo measurements observe the UP-DOWN states by building accumulative excitatory recurrences that are controlled through a downscaling process of synaptic weight (Bartram et al., 2017). Transitions in the sleep-cycle have also been modelled through the interactions of neurotransmitters that modulated hyperpolarization and depolarization of each neuron (Krishnan et al., 2016). The cumulative control of neuromodulation between these mechanisms yields the overall effect of a self-sustaining UP-DOWN regime, or an asynchronous noisy regime. In this work, the specific pathway by which this is achieved is abstracted to the global adaptiveness that is tuned to the state of each region. What is crucial is that the capacity to adapt to increased excitatory inputs allows the HDMF model to enter a slow-wave regime that is not accessible in the standard formulation. Thus, the gain of embedding this plasticity rule in the DMF is not only homeostatic control, but also an expansion of the model’s dynamical repertoire toward bistable UP–DOWN activity.

### Coexisting Dynamical States and Chimeric-like Activity

The HDMF generated distinct dynamical regimes simultaneously, including UP–DOWN states and asynchronous activity. This feature resonates with empirical observations of local sleep (Andrillon & Oudiette, 2023), partial awakenings, and dissociative conditions(Antelmi et al., 2016), where different brain areas can manifest oscillatory sleep-like states that concur with asynchronous activity (Stasinski et al., 2024). When this happens in the HDMF we refer to it as a “chimeric-like” state. In complex systems, a “chimeric system” exhibits different dynamics across identical regions(Haugland, 2021). Although this is not the case for our model –regions have different connectivity patterns and different learning rates–, it is neither the case for the brain. More broadly, several modeling approaches highlight that embedding empirical heterogeneity is a fruitful direction for capturing realistic dynamics (Patow et al., 2024; Perl et al., 2023). In the HDMF heterogeneity in adaptation gave rise to different dynamics that not only co-existed but influenced each other, and interacted effectively via structural coupling. This capacity could help illuminate the role of SWA within asynchronous regimes, the use-dependent regional changes in homeostatic regimes^36^ and potentially lesioned areas of the brain. Additionally it may explain how certain states (e.g., lucid dreams, false awakenings) could emerge through specific pathways that shift local inhibition and excitatory drive (Aru et al., 2020; LaBerge et al., 2018).

### Performance of the HDMF Relative to Prior DMF Applications

We fitted both the DMF and HDMF to resting-state fMRI data. Different observables were optimally fitted in distinct regions of the explored parameter space. For a fair comparison, we also included DMF parameter points that are typically excluded due to excessively high firing rates. Although it has been shown that not all observables can be optimally fitted at a single working point in the DMF (Luppi et al., 2025), we chose to compare the models in regions where both FC and FCD were reproduced at an acceptable level. The HDMF showed comparable performance for FC and improved fitting to FCD relative to the DMF in the current dataset. Importantly, the overlap regions yielding acceptable FC and FCD fits did not colocalize with the regions expressing sleep-like slow-wave activity, consistent with the wake-like nature of the resting-state empirical data. This improvement in the fit to FCD suggests that the homeostatic mechanism enhances the model’s ability to capture empirical dynamical structure, together with the previously described increase in robustness and dynamical repertoire.

The HDMF remained stable under neuromodulation across a range of intensities, whereas the static FIC often failed to maintain firing rates within a biologically plausible regime. This was reflected in the strong correlation between regional receptor density and resulting activity in the HDMF. In contrast, the DMF and the mixed model—where the FIC is derived from the average of the dynamic FIC—exhibited weaker correlations and pronounced hyperexcitability. Importantly, this increased stability did not come at the cost of empirical fitting performance. When reproducing the LSD fitting analysis of Herzog et al., the HDMF maintained firing rates within a narrower biologically plausible range and provided a better fit to empirical pharmacological fMRI data than the DMF. The ability to preserve stable dynamics while improving the fit to receptor-specific pharmacological perturbations is particularly relevant for modeling approaches aimed at developing digital twins to test pharmacological strategies (Alnagger et al., 2026; Mindlin et al., 2024)

### Comparison to Other Model Families

The plasticity rule we implement is biologically inspired by how inhibitory synapses regulate the balance between excitation and inhibition, following the work by Vogels et al. Other studies have also implemented it for large-scale networks. Stasinski and colleagues (Stasinski et al., 2024) show that by incorporating it there is a complexity increase in local activity, as well as a more constrained and accurate space for optimal fitting of BOLD data. In their framework the dynamic FIC variable is mainly used as a tuning mechanism: after convergence, the final dynamic values are averaged to obtain static regional inhibitory parameters for post-FIC simulations. In another study (Hellyer et al., 2016), the same was done with the Wilson-Cowan model, showing that FIC adaptation to specific target rates can yield power-law distributed avalanche sizes. This suggested that the homeostatic system was at criticality, explaining its enhanced flexibility. This class of mechanism should be distinguished from other forms of adaptation used in large-scale models. For example, AdEx-based models (Goldman et al., 2023) implement spike-frequency adaptation, in which an intrinsic adaptation current accumulates with excitatory firing and decays during periods of low activity. This form of adaptation can also generate Up–Down states, but it is mechanistically different from the Vogels-type inhibitory plasticity rule. Rather than proposing a new plasticity rule for the modeling literature, this study focuses on the implications of tuning this established rule to achieve homeostasis in the DMF model. Specifically, we modified the aforementioned rule by adding a *τ*_decay_ parameter, which was necessary to reach homeostasis (Supplementary Figure 2). This decision changes the rule from the accumulation of activity towards a target rate to an on-going process that needs to be kept alive, making it more akin to a homeostatic process. Furthermore, the Frequency-Current curve of the DMF generates firing rates from currents through an exponential-linear function that does not plateau, i.e. can increase indefinitely, contrary to typical sigmoid functions used in other models, such as the Wilson-Cowan. The *τ*_decay_ works as an adaptive threshold that can promote potentiation or depression according to the current state, penalizing the FIC in a magnitude-dependent way (the larger the FIC, the larger the penalization), which in turn stabilizes the activity without imposing a saturation point. It is worth noticing that in contrast to other models that externally impose saturation through a sigmoid, in the HDMF a comparable constraint emerged endogenously through the adaptive threshold. In this sense, the regulation of activity was not hardwired but self-organized within the system’s dynamics. This control mechanism prevents runaway excitation characteristic of pathological states such as epilepsy.

### Broader Applications and Future Directions

Because the HDMF can generate slow-wave activity, it also provides a framework to introduce explicit temporal structure into adaptive processes. This opens the possibility of modeling transient, spatially localized slow-wave events resembling “local sleep” during otherwise wake-like activity (Andrillon et al., 2021). When combined with probe-based resting-state paradigms, such as experience sampling during mind wandering (Mortaheb et al., 2022), this framework could be used to test whether specific temporal rules of adaptation align with moments of mind blanking or attentional disengagement.

In addition, we showed that the HDMF can accommodate neuromodulatory influences as external perturbations acting on internal network dynamics. This opens the possibility of modelling transient pharmacological effects in the DMF, such as the short-lasting changes induced by DMT during fMRI acquisition. In this setting, the time course of the drug can be introduced as a continuous change in neuromodulatory gain, mimicking both the intensity of the psychedelic effect and its decay over the fMRI acquisition (Piccinini et al., 2025), while the dynamic FIC preserves firing-rate stability throughout the perturbation. This approach can be extended to other forms of extrinsic input, including auditory or visual stimulation (Bekinschtein et al., 2009; Türker et al., 2023). In this context, the HDMF provides a setting in which slow resting-state fluctuations coexist with adaptive mechanisms that regulate responses to external information, allowing resting-state dynamics and task-related processing to be studied within the same model.

Beyond reproducing a diverse set of brain states, homeostatic inhibitory control in the HDMF could prove valuable in personalized connectome-based modeling, where individual structural or PET receptor maps may push local network dynamics into regimes of pathological hyper- or hypo-activity (Maestú et al., 2021; Ranasinghe et al., 2022), or cortical development (Zhang et al., 2024). Adding region-specific target firing rates or multi-timescale plasticity might further enhance the model’s ability to simulate neurological disorders –such as epilepsy, brain lesions or neurodegenerative conditions– and to explore therapeutic strategies. Combining homeostatic and Hebbian-like rules could help capture learning-dependent processes alongside baseline stability.

### Limitations

Although the HDMF addresses important hurdles in large-scale brain modeling, several limitations remain. First, our approach focuses on a single target firing rate, which may not reflect the multiple concurrent homeostatic targets in the brain –such as metabolic constraints or different firing rates across regions. To deepen the biological sense of homeostasis, these processes should be added, since they may affect firing-rate but are not necessarily through their produced outcome. Also, the LR-*τ*_decay_ relationship needs to be obtained for every combination of structural connectivity and target firing rate, as shown by our analytical derivation. However, this can be done in a comparatively shorter time than the original pseudo-gradient algorithm (which also lacks the FIC dynamics of the HDMF). Second, we tested primarily cortical connectivity, leaving open questions about subcortical and thalamocortical contributions. Third, we did not explore interactions between homeostatic plasticity and other plasticity mechanisms (e.g., Hebbian or STDP), which could be critical for understanding learning and memory formation. Nevertheless, learning and memory may occur at much finer spatial scales than the ones modelled with whole-brain models, making these limitations transversal to the field of whole-brain modelling. Finally, further empirical validation, including clinical populations or perturbation experiments, will be necessary to confirm the predictive utility of the HDMF across diverse real-world contexts.

## Conclusions

By adding a homeostatic plasticity rule to the DMF model we showed how this mechanism is beneficial for ensuring stability and enhancing realism. This also allowed the modelisation of slow waves, which is unprecedented in the DMF model. We finally propose that by adding heterogeneity in homeostatic properties we can recreate chimeric-like states, suggesting a potential mechanism for sleep dissociative states. Altogether, the HDMF illustrates how embedding homeostatic self-regulation into whole-brain models can overcome fragility, extend dynamical repertoire, and provide mechanistic links to complex brain states.

## Materials and methods

### Participants

#### fMRI data

Empirical resting-state fMRI data were obtained from the open processed HCP-YA functional connectome dataset(*Derived Products from HCP-YA fMRI*, 2022), derived from the Human Connectome Project Young Adult S1200 release (Van Essen et al., 2012). We used the no-global-signal-regression version of the resting-state BOLD time series parcellated into 100 cortical regions using the Schaefer atlas. The original HCP-YA dataset includes 3T MRI and behavioural data from healthy young adults aged 22–37 years, with resting-state fMRI acquired at TR = 0.72 s and 1200 time points per run. We restricted the analysis to the 99 uncorrelated subjects (Afyouni & Nichols, 2018).

The data used here had already undergone the HCP minimal preprocessing pipeline (Glasser et al., 2013). Briefly, this includes correction of spatial distortions, motion correction, bias-field correction, registration to standard space, mapping to CIFTI grayordinate space, and surface/parcel-constrained smoothing. For resting-state fMRI, the HCP-provided data also include 24-parameter motion regression and ICA-FIX denoising. The processed dataset further provides versions with and without global signal regression; we used the version without global signal regression. Resting-state time series in the processed dataset were band-pass filtered using a 2nd-order Butterworth filter between 0.01 and 0.1 Hz. Functional connectivity was computed as the Pearson correlation matrix between all pairs of regional BOLD time series. Functional connectivity dynamics were estimated using a sliding-window approach, with a window length of 84 TRs and a step size of 4 TRs. The resulting windowed FC matrices were used to compute the empirical FCD structure for model fitting.

#### Structural connectivity for empirical analysis

Empirical structural connectivity was obtained from the publicly available multimodal connectivity dataset released by Hansen et al (Hansen et al., 2023). The structural connectomes were derived from diffusion-weighted imaging data from 326 unrelated HCP participants from the S900 release (Van Essen et al., 2013). The source data were preprocessed with MRtrix3: fiber orientation distributions were estimated using multi-shell multi-tissue constrained spherical deconvolution, white-matter streamlines were reconstructed with probabilistic tractography, and streamline weights were optimized using cross-section multipliers. Hansen et al. constructed a group-consensus structural network designed to preserve the density and edge-length distributions of individual connectomes. Edge weights were assigned by averaging the log-transformed streamline counts of nonzero edges across participants and were subsequently rescaled between 0 and 1. The resulting connectivity matrices are openly available through the authors’ GitHub repository. In the present study, we used the Schaefer-100 version of the group-consensus structural connectivity matrix, matching the cortical parcellation used for the empirical fMRI data. No additional tractography or diffusion MRI preprocessing was performed. For simulations of empirical fMRI data, the SC was normalized by its maximum eigenvalue.

#### Structural connectivity enrichment

To improve the agreement between structural and empirical functional connectivity, we enriched the SC matrix following a model-based iterative procedure proposed by previous work (Deco, McIntosh, et al., 2014; Ipiña et al., 2020) on optimal structural connectivity estimation. Enrichment was performed only with the standard DMF model, using fixed parameters G=1 and α=0.7. At each iteration, simulated FC was compared with empirical FC of the respective groups consensus of Hansen et al (Hansen et al., 2023), and all nonzero SC weights were updated according to:

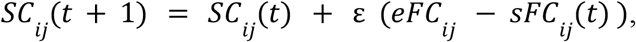

where *eFC*_*ij*_ and *sFC*_*ij*_ denote empirical and simulated FC, respectively, and *ε* is the enrichment step size (0.01). The update was restricted to nonzero entries of the original group-consensus SC matrix, preserving its binary topology and introducing no new structural edges.

#### Structural connectivity data for theoretical analysis

Diffusion-weighted MRI data were obtained from a previously collected cohort of 16 healthy right-handed participants (5 women; mean age ± s.d., 24.8 ± 2.5 years) scanned at Aarhus University, Denmark (Deco et al., 2017). The study protocol was approved by the local ethics committees, and all participants provided written informed consent. Imaging was performed in a single session on a 3 T Siemens Skyra scanner. Diffusion data were acquired with 62 nonlinear diffusion-encoding directions (b = 1500 s/mm^2^), interleaved with non-diffusion-weighted images (b = 0), using a repetition time of 9000 ms and an echo time of 84 ms. Images were collected with a voxel size of 1.98 × 1.98 × 2 mm^3^ and reconstructed on a 106 × 106 matrix, with both anterior-to-posterior and posterior-to-anterior phase-encoding directions. Motion and eddy-current correction was performed using MRtrix (Tournier et al., 2019). Diffusion images were co-registered to each participant’s T1-weighted anatomical image and then to MNI space. The AAL atlas was then transformed from MNI space to individual native diffusion space using the inverse transformations, preserving discrete labels via nearest-neighbour interpolation. Fibre orientation distributions were estimated using constrained spherical deconvolution, and whole-brain probabilistic tractography was performed to reconstruct inter-regional pathways. Streamlines were aggregated between atlas-defined regions to generate individual structural connectivity matrices, which were subsequently averaged across participants to obtain a group-representative connectome Further details are provided in the original study (Deco et al., 2017).

#### PET-derived receptor density data

We incorporated publicly available positron emission tomography (PET) receptor density maps from the normative neurotransmitter atlas published by Hansen and colleagues (Hansen et al., 2022). These maps include whole-brain estimates of multiple neurotransmitter receptor systems aggregated across healthy individuals, allowing in-vivo quantification of receptor distributions. For the serotonin 2A receptor (5-HT_2_A) specifically, the atlas is based on PET imaging using the agonist radioligand Cimbi-36, which binds with high affinity to 5-HT_2_A receptors and has been validated as a marker of their density in the living human brain. PET tracer maps were registered to MNI space using the Neuromaps (Markello et al., 2022) library and parcellated using AAL atlas to extract mean receptor density values for each region denoted as *D*_*i*_. Density values were normalised by dividing by the maximum value, ensuring that *max*(*D*_*i*_) = 1.

### Dynamic mean field (DMF) model

We used the DMF to simulate the brain activity of each brain region, as proposed in (Deco, Ponce-Alvarez, et al., 2014b). The DMF simulates whole-brain activity by defining the local dynamics of each region as two interconnected pools of excitatory (E) and inhibitory (I) neurons while long range interactions are only between the E pools of each region, adjusted by the structural connectivity *C*_*ij*_ (for details, refer to Methods - Anatomical Connectivity).

The model is based on the earlier work of (Wong & Wang, 2006), where excitatory synaptic currents 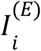 are mediated by NMDA receptors, while inhibitory currents 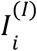 are mediated by GABA receptors.

In detail, the DMF is defined by the following system of coupled differential equations:

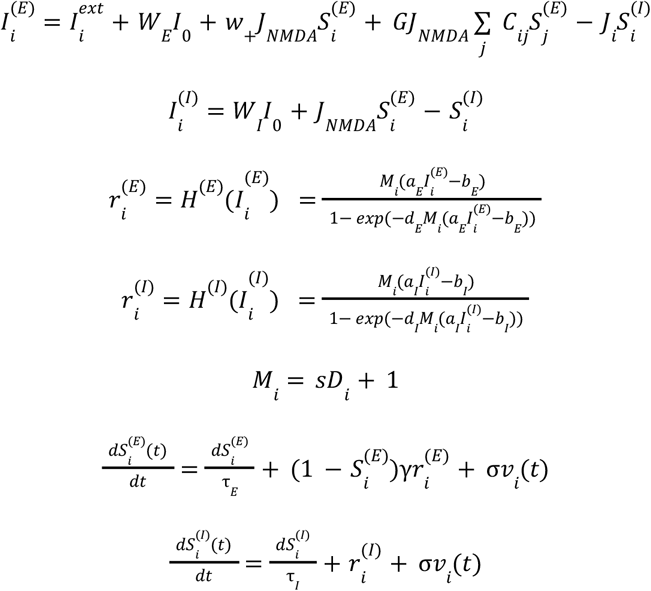

Here, 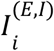 (in nA) represents the total input current, and 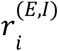 (in Hz) represents the respective firing rates. 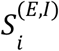 represent the synaptic gating variables of each pool. The neuronal nonlinear response functions, *H*^(*E,I*)^ , convert the total input currents received by the E and I pools into firing rates, 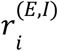, based on the input-output function proposed by (Abbott & Chance, 2005). *H*^(*E,I*)^ is also known as the F-I curve. The model employed specific gain factors, threshold currents, and constants to determine the shape of the curvature of *H*^(*E,I*)^ around the threshold. *s* is called the neuromodulatory gain, which scales the impact of the receptor density *D*_*i*_ in each region, which is mapped into *M*_*i*_ (Deco et al., 2018). G is a free parameter called the global coupling, which scales the influence of the network on each region. See Table 1 for the name and value of each model parameter.

**Table 1.**
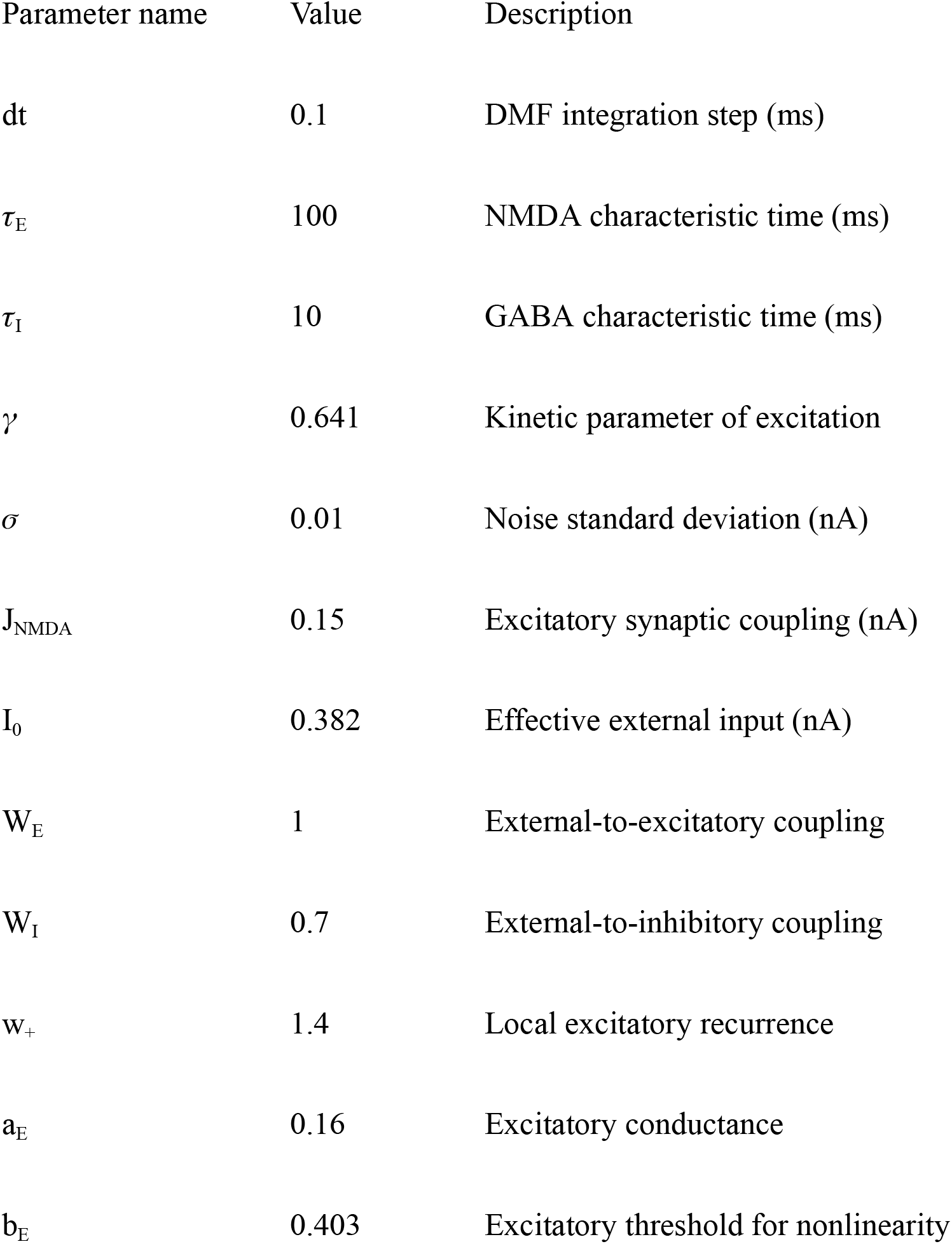
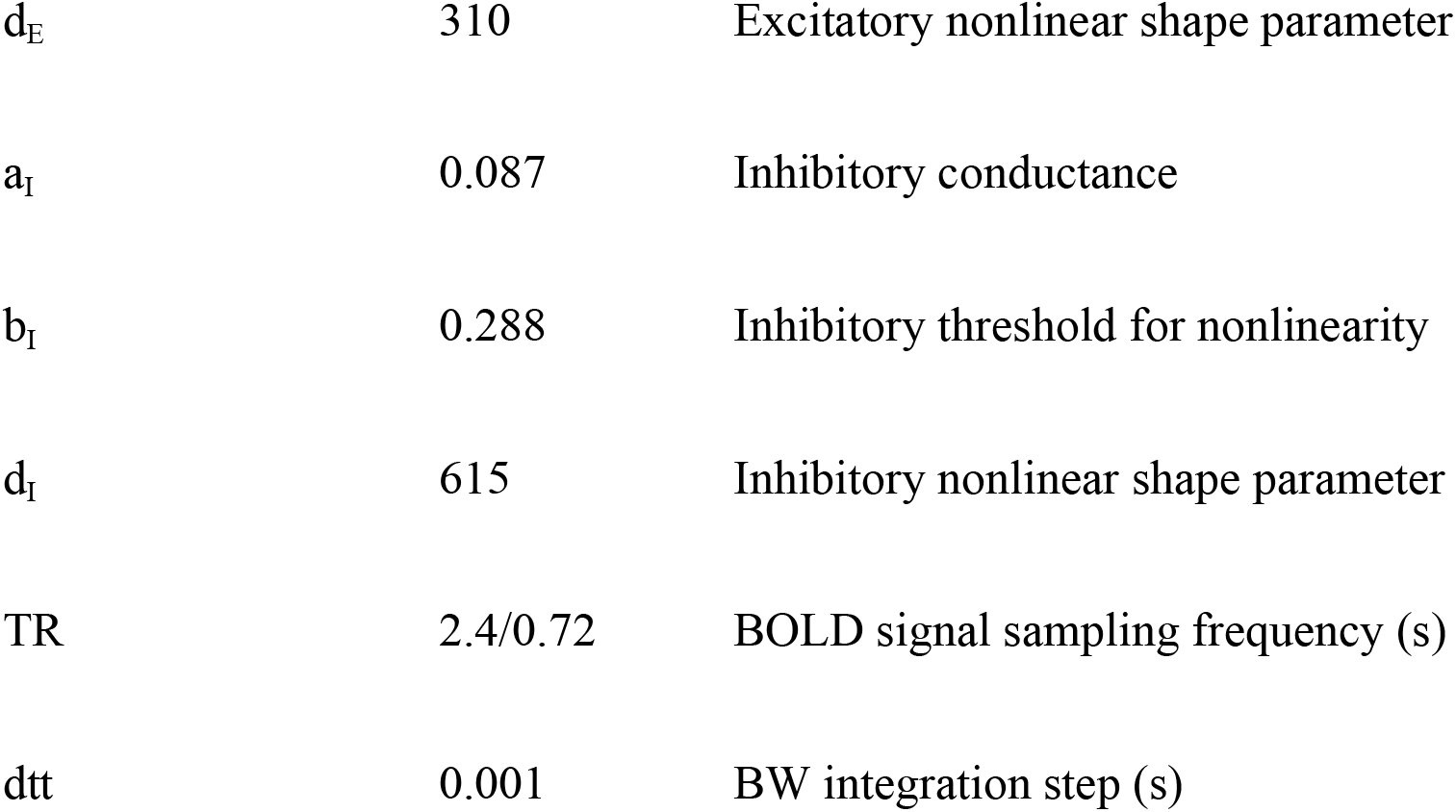
Model parameters.

### Feedback inhibitory control (FIC)

The parameter *J*_*i*_ , referred to as FIC, defines the synaptic weight from the inhibitory (I) to the excitatory (E) population. As demonstrated by Deco and colleagues(Deco, Ponce-Alvarez, et al., 2014a) precise tuning of this parameter allows the excitatory firing rates to be clamped around 3.4 Hz, maintaining them within a biologically plausible regime. Because *J*_*i*_ must be optimized for each value of the global coupling parameter G, this step substantially increases the computational cost of the simulations. However, Herzog et al (Herzog, Mediano, Rosas, Luppi, Sanz Perl, et al., 2024) proposed a heuristic linear solution for the FIC which derives the optimal value from a linear relationship between G and the anatomical connectivity strength β as follows:

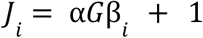

where α is a free parameter –representing the global E/I balance– that depends on the parcellation and in the target firing rate, and 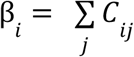 , i.e. the anatomical connectivity strength. This solution achieves stability of firing rates up to critical value of G, where this solution cannot control the firing rates anymore and the system enters into a hyperactivity regime. See (Herzog, Mediano, Rosas, Luppi, Sanz Perl, et al., 2024) for a detailed description of this approximation.

### Inhibitory plasticity rule with decay

Following previous work by Hellyer and colleagues (Hellyer et al., 2016), we implemented a local inhibitory plasticity rule to dynamically adapt *J*_*i*_ such that the excitatory average firing rates converge to a target firing rate ρ. We extended this plasticity rule to include a decay term τ_*decay*_ on the synaptic weight. This rule follows:

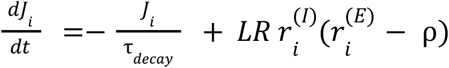

LR is the learning rate and represent the speed of the update of the synaptic weight by an active mechanisms while τ_*decay*_ represents an adaptive threshold mechanism: when excitatory firing rates exceed the target above a threshold, inhibition will be potentiated; and vice versa, firing rates below the threshold will induce a depression in inhibition.

### Homeostatic dynamic mean field model (HDMF)

The inhibitory plasticity rule enables a dynamic control of the local inhibition based on the relationship between the E and I pool and the excess or lack of excitatory activity. However, for some combination of parameters the average excitatory firing rate will not converge to the target, but will converge to some other value, failing to achieve homeostasis. To fully achieve homeostasis, i.e. that the average firing rates converge to the expected target, we used the following heuristic relationship between the LR and τ_*decay*_:

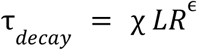

Then, τ_*decay*_ has a power-law relationship with LR controlled by the coefficient χ and the exponent ϵ, which are obtained by numerical simulations (see Figure 2A). This implies that τ_*decay*_ depends on LR and then the plasticity rule can be fully expressed in terms of LR. As shown in Figure 2A, these parameters depend on the target firing rate but, regardless of this target, this relationship achieves maximal homeostasis in a wide range of G values, i.e. minimizes the difference between the target and the average firing rates. To distinguish the DMF from its version with homeostasis, we call the latter homeostatic DMF (HDMF). For the empirical analysis we recalculated the necessary homeostatic coefficients for the used connectivity weights. This is explained in Supplementary Figure 6.

All time variables in the neural model are expressed in simulated biological time. The stochastic differential equations were integrated with a time step of dt = 0.1 ms. The decay constant should be interpreted as the relaxation time scale of the effective inhibitory feedback variable. The range explored here spans sub-second to tens-of-seconds dynamics, which is compatible with fast adaptive regulatory processes in neural systems, without implying a one-to-one correspondence with a specific molecular mechanism.

#### Analytical derivation of the LR-τdecay power-law relationship

In stationarity, the plasticity rule for the local inhibitory weight *J*_*i*_ admits a closed-form relation between the learning rate *LR* and the τ_*decay*_. Writing 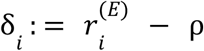 for the excitatory deviation from the target, the homeostatic regime is characterised by E [δ_*i*_] = 0, i.e. the expected deviation from the target is zero with non-zero variance. Then, by taking expectations over time we obtain:

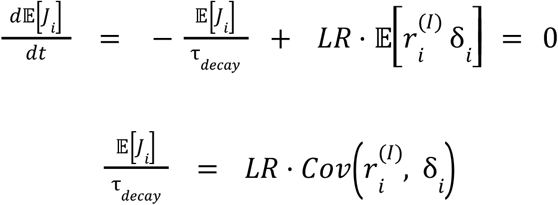

where E [δ_*i*_]= 0 has been used to identify the right-hand side as a covariance. Rearranging,

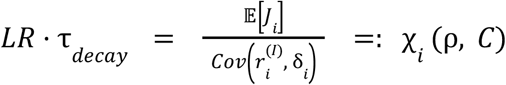

This is the central result: the product *LR* · τ_*decay*_ is fixed by stationary network statistics, so τ_*decay*_ ∝ *LR*^−1^. The prefactor χ_*i*_ (ρ, *C*) is set entirely by E[*J*_*i*_ ] and 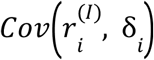. Note that the excitatory deviation *δ*_*i*_ is, by definition, the displacement of 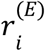 from the target ρ; its statistics therefore depend on ρ directly. Also, the local inhibitory rate 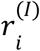 is driven by the excitatory pool, which in turn receives the inputs from the network shaped by the connectivity matrix C. Thus, its co-variation with *δ*_*i*_ depends on *C*. The prefactor is therefore non-universal: it must be recalibrated for each combination of ρ and *C*.

The canonical scaling *LR*_*decay*_ ∝ τ ^−1^ is the special case in which the right-hand side of the master equation is independent of τ_*decay*_. To quantify deviations from this case, the relevant observable is the exponent of the power law itself, i.e. the slope of *LR* against τ_*decay*_ in log–log coordinates,

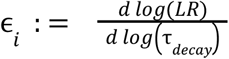

which equals − 1 in the canonical case and departs from it otherwise. To extract this slope from the master equation, we take logarithms,

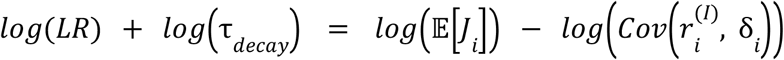

and differentiate with respect to *log* (τ _*decay*_):

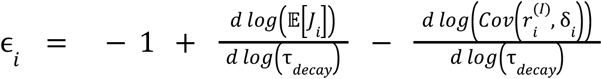

The canonical value ϵ_*i*_ = − 1 is recovered when both E[*J*_*i*_ ] and 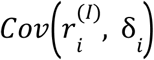 are independent of τ_*decay*_ ; any dependence translates one-to-one into a shift of the exponent. The deviation of ϵ_*i*_ from − 1 is therefore itself a function of ρ and *C*; like the prefactor, the effective exponent must be recalibrated whenever either is changed.

### Simulating BOLD signals

To convert the simulated mean field activity into a BOLD signal, we used the generalized hemodynamic model by Stephan et al (Stephan et al., 2007). The BOLD signal for each brain region *i* was calculated based on the firing rate of the excitatory pools, 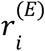. In this framework, an increase in firing rate caused a rise in a vasodilatory signal, *s*_*i*_ , which was also subject to auto-regulatory feedback. Consequently, changes in blood inflow, *f*_*i*_ , were influenced by this signal, leading to variations in blood volume, *v*_*i*_ and deoxyhemoglobin content, *q*_*i*_. Also, for sustained firing increase, bold response saturates (Rosa et al., 2011; Tesler et al., 2023). We modeled this neurovascular coupling with baseline invariant sigmoid *g*. The interactions between these biophysical variables were described by the following equations:

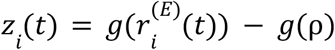

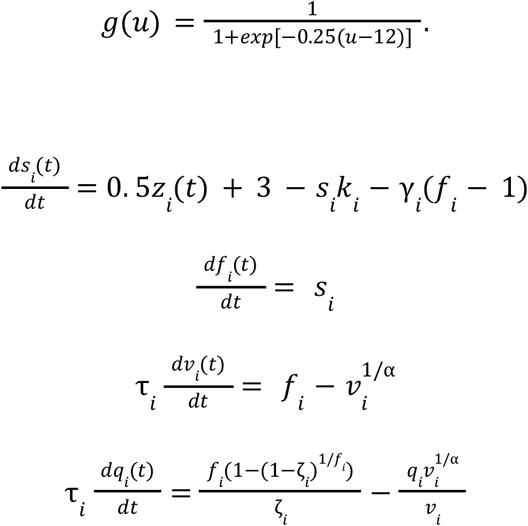

where ζ is the resting oxygen extraction fraction, τ is a time constant and α that represents the resistance of the veins. Finally, the BOLD signal in each area *i,B*_*i*_ , is a static nonlinear function of volume, *v*_*i*_ , and deoxyhemoglobin content, *q*_*i*_ , that comprises a volume-weighted sum of extra- and intravascular signals:

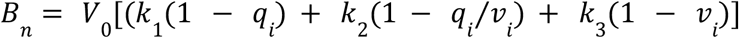

All biophysical parameters were adopted from reference (Stephan et al., 2007). To isolate the frequency range most relevant for resting-state activity, both empirical and simulated BOLD signals were band-pass filtered between 0.01 and 0.1 Hz.

### Bayesian optimization of target firing rates

We used Bayesian optimization, implemented with Optuna’s Tree-structured Parzen Estimator (TPE) sampler, to fit a region-specific target firing-rate vector for HDMF. TPE is a sequential, model-based search strategy that builds separate density estimates for promising and less promising regions of parameter space based on previously evaluated trials. New candidates are then proposed by favoring parameter values that are more likely under the “good” density than under the “bad” one, allowing efficient exploration of high-dimensional, non-convex spaces without requiring gradients. The search was initialized using homogenous target = 3.44 for all regions and kept the previously identified optimal global coupling and learning rate fixed (G = 1.1, LR = 0.8), with the decay constant derived from the calibrated LR–decay relationship. Target was constrained to the [1,5] range and the LR–decay relationship was not re-calibrated. For each candidate, we evaluated model performance using the same fixed seed set and preprocessing pipeline as in the homogeneous model, and defined the objective as the Pearson correlation between the population-averaged simulated FC and the population-averaged empirical FC. Optimization stopped after 100 iterations and the best solution was used.

### Fit to empirical data

Model parameters were optimized selecting optimal points in an exploratory grid. The objective function was defined either as: (i) the Pearson correlation between the simulated and population-averaged empirical FC matrices, or (ii) the Kolmogorov–Smirnov (KS) distance between the histogram of the pooled empirical Functional Connectivity Dynamics (FCD) and that of the simulated FCD. Each simulation generated 600 seconds of BOLD signals sampled at TR = 0.72 s, yielding a number of time points comparable to the empirical data. The optimization assumed a stochastic objective function, allowing the algorithm to randomly initialize each simulation and estimate variability due to different initial conditions.

### Functional connectivity and functional connectivity dynamics

Functional connectivity (FC) was estimated by computing the Pearson correlation between the empirical (simulated) fMRI time series. For the functional connectivity dynamics (FCD) we used the same approach, but we computed the FC in a sliding window of 84 TR and step size of 4TR. Then, the upper triangular part of the FC of each window was transformed into a vector and the Pearson correlation between all the time windows was computed, obtaining the FCD matrix. This matrix captures the correlation of FC patterns across time.

### Detection of oscillatory dynamics

We computed the power spectral density (PSD) of neuronal firing-rate traces using Welch’s method with a 1000 Hz sampling frequency, segment lengths of 4000 samples, and a 50% overlap. Each PSD was evaluated only up to 100 Hz, and the maximum point of this spectral region was identified to determine the dominant oscillation frequency and its corresponding power. This approach produced four principal outputs: the distribution of frequencies (freqs), the PSD values for each frequency (psd), the peak frequency (max_freqs), and the maximal PSD within 0–100 Hz (max_power), providing a concise characterization of each signal’s strongest oscillatory component.

#### Computing regime type index

To quantify how strongly a network endowed with region‐specific plasticity deviated from either of the two baseline uniform regimes, we defined the metric “Regime type index” (RTI) that yields a value for each region. For each analysis we first identified a subset of *N* cortical regions that were assigned a learning‐rate parameter LR_HET_ while all remaining areas retained the reference uniform value LR_UNI_. One hundred independent simulations were then run with this heterogeneous vector. For every run we extracted the dominant oscillation frequency of each region (see Detection of oscillatory dynamics) and averaged these frequencies across simulations, yielding 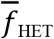. Each region uses two reference values obtained from fully uniform networks 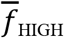 from the “asynchronous” high‐learning‐rate regime and 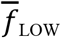 from the “slow‐wave–like” low‐learning‐rate regime. The RTI rescales the position of 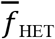 between these anchors according to

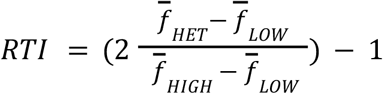

With this normalization the index equals –1 when the heterogeneous network is indistinguishable from the low‐learning‐rate regime and +1 when it fully reproduces the high‐learning‐rate regime, while intermediate values indicate partial convergence toward either regime.

### Statistics

The distributions of goodness-of-fit values for the FC and FCD histograms were compared using Kolmogorov–Smirnov test.

## Acknowledgments

We thank the financial support of the FLAG-ERA research funding organisation (project ModelDXConsciousness). The collaboration between RH and IM was supported by the María de Maeztu Program for units of Excellence in R&D, grant CEX2021-001164-M/10.13039/501100011033.

## Author contributions

Conceptualization: Iván Mindlin, Rubén Herzog

Methodology: Iván Mindlin, Rubén Herzog, Andrea Luppi, Carlos Coronel, Marc Llabrés

Investigation: Iván Mindlin, Rubén Herzog

Visualization: Iván Mindlin Supervision: Rubén Herzog

Writing-original draft: Iván Mindlin, Rubén Herzog

Writing-review & editing: Iván Mindlin, Carlos Coronel, Jacobo D. Sitt, Rodrigo Cofré, Andrea Luppi, Thomas Andrillon, Yonatan Sanz Perl and Rubén Herzog

## Data and materials availability

Computational codes to run the HDMF and reproduce the results are available at https://github.com/Picardian14/fastHDMF-code.git. We’ve also made a pure Python implementation of the same model. https://github.com/carlosmig/Homo_DMF.

