## Supplementary Figures for "The impact of homeostatic inhibitory plasticity in a generative biophysical model"

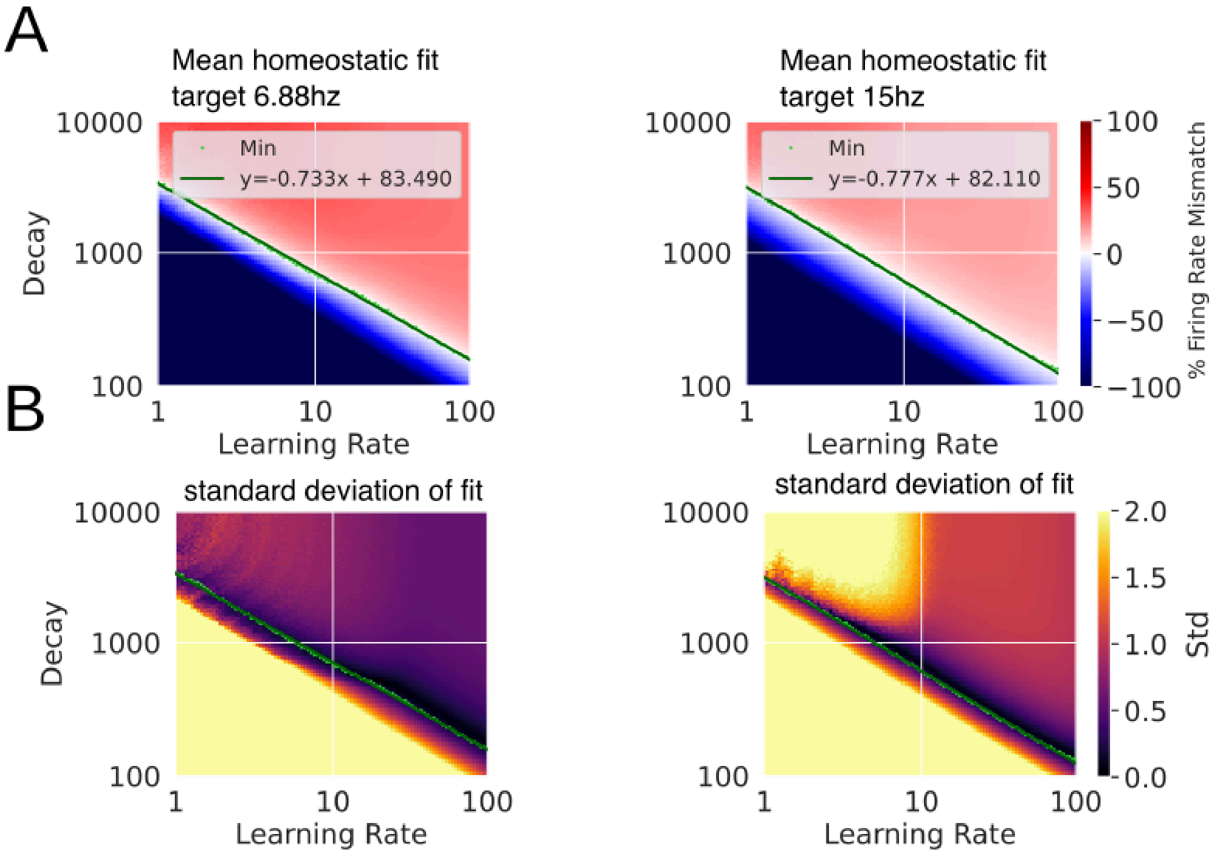


**Supplementary Figure 1.** Similar to Figure 2, but for additional target rates. The HDMF can achieve homeostasis to different firing rates with a linear fit. **A** Average percentage of mismatch between the average E firing rate and ρ **B** Same as in **A**, but showing the standard deviation in the same range of G values.

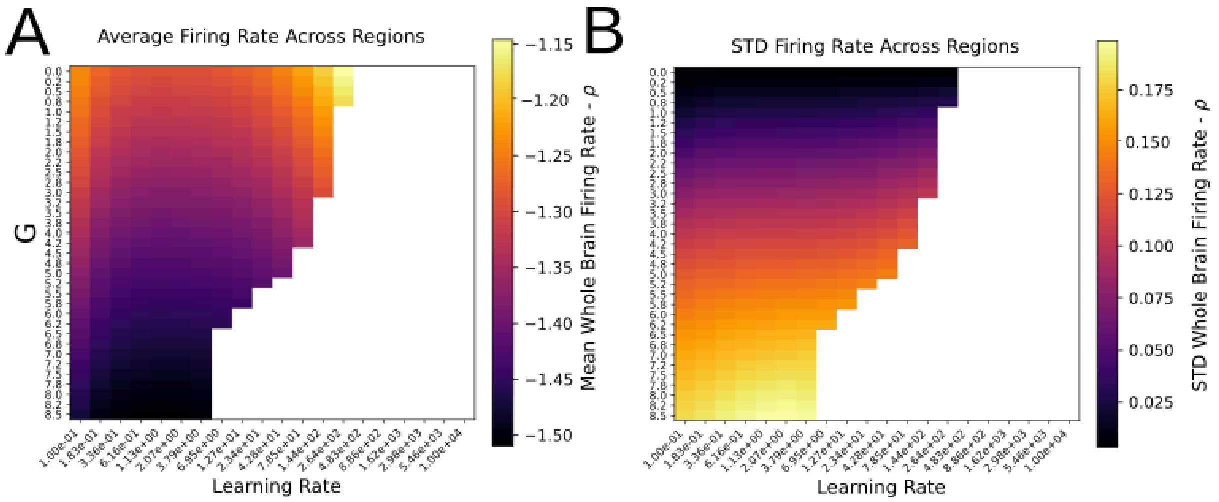


**Supplementary Figure 2.** Mean (**A**) and standard deviation (**B**) homeostatic fit for plasticity rule without decay. Empty values represent exploding firing rates.

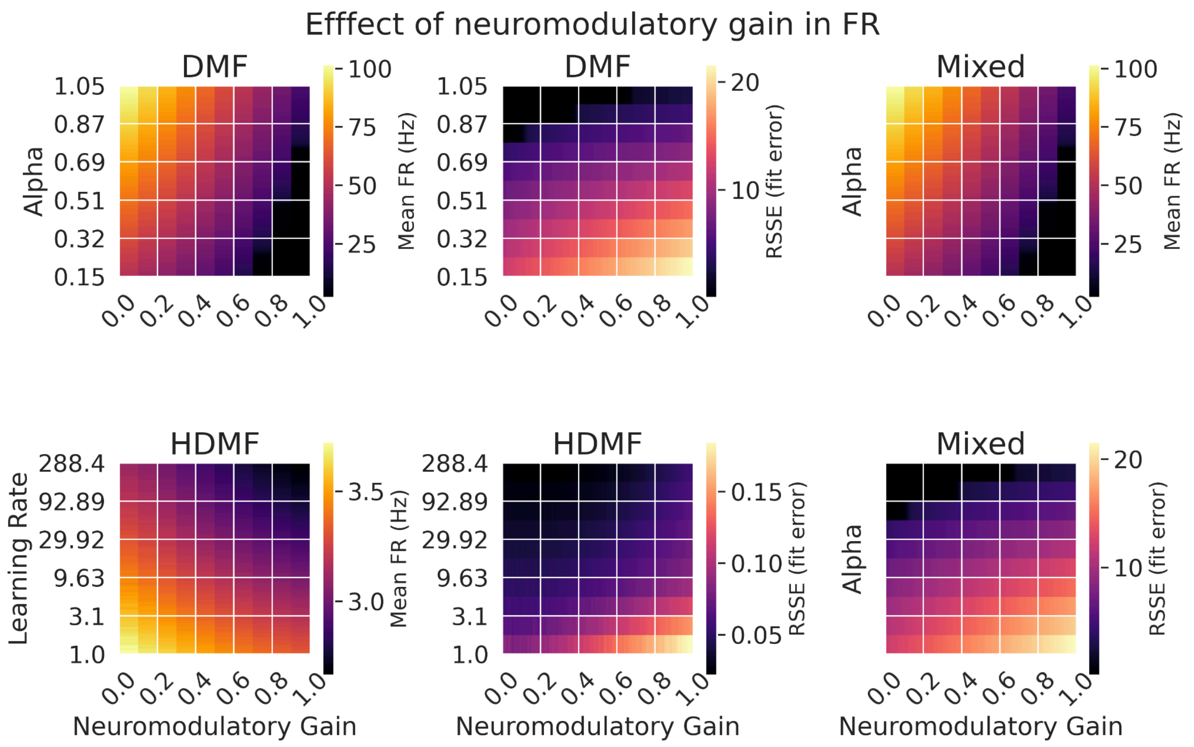


**Supplementary Figure 3.** Generalization of the tendency shown in Figure 3. in the manuscript. **A-D** Show the average firing rate across all regions for each pair of parameters of the DMF and HDMF respectively **C** Similar, but since there is only neuromodulation we show the variability across repetitions. **B-E-F** Show the RSSE for the linear fit of receptor density with respect to firing rates for each region.

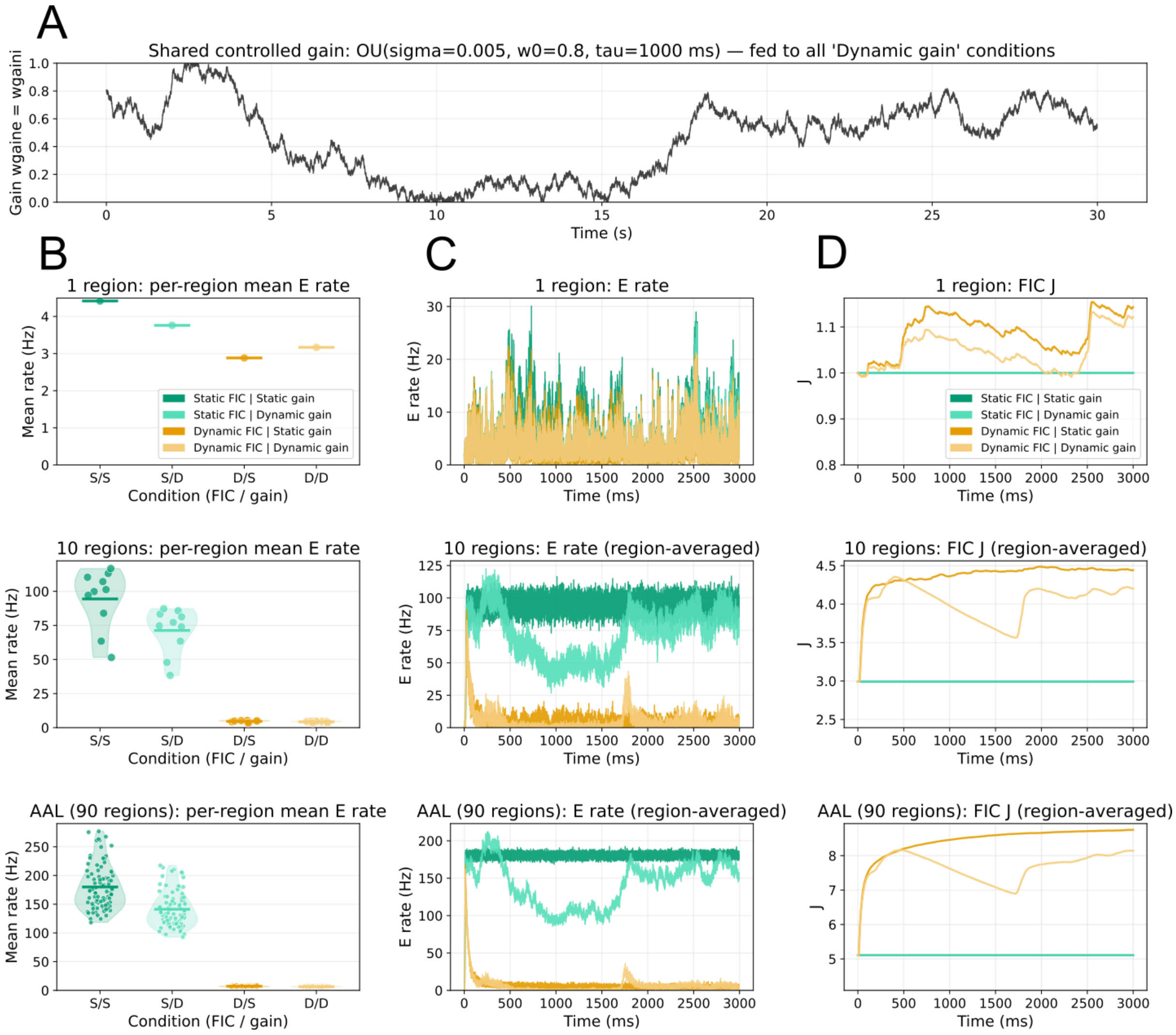


**Supplementary Figure 4**. **Interactions between dynamic gain and feedback inhibition control across network scales.** (A): Shared gain trajectory used across all dynamic-gain conditions, sampled from an Ornstein-Uhlenbeck process ($\sigma = 0.005$, $w_0 = 0.8$, $\tau = 1000$ ms, clipped to [0, 1]). Gain modulation was applied using the same 5-HT2A PET receptor density map shown in Figure 3. For static-gain conditions, the gain value was fixed to the temporal mean of the dynamic gain trajectory shown in (A). The dynamic gain trajectory in (A) was used in all conditions presented in panels (B–D). (B–D): Three network configurations (1 region, 10 regions, AAL 90 regions) evaluated under four conditions combining static FIC (DMF) dynamic FIC (Homeostatic DMF, HDMF) and static/dynamic gain. Within each network size, columns present (B) per-region mean excitatory rates with individual data points and strip-violin distributions, (C) excitatory firing rate time series (single trace for $N = 1$; regional mean for $N \geq 10$), and (D) feedback inhibition control coefficient J (single trace for $N = 1$; regional mean for $N \geq 10$). Green colors denote DMF; yellow denotes HDMF. Darker tones indicate static gain; lighter tones indicate dynamic gain (DMF/S, DMF/D, HDMF/S, HDMF/D). The HDMF maintains firing rates close to the target regime despite abrupt gain fluctuations, whereas DMF transitions into a sustained overexcited state under both dynamic gain and static neuromodulation.

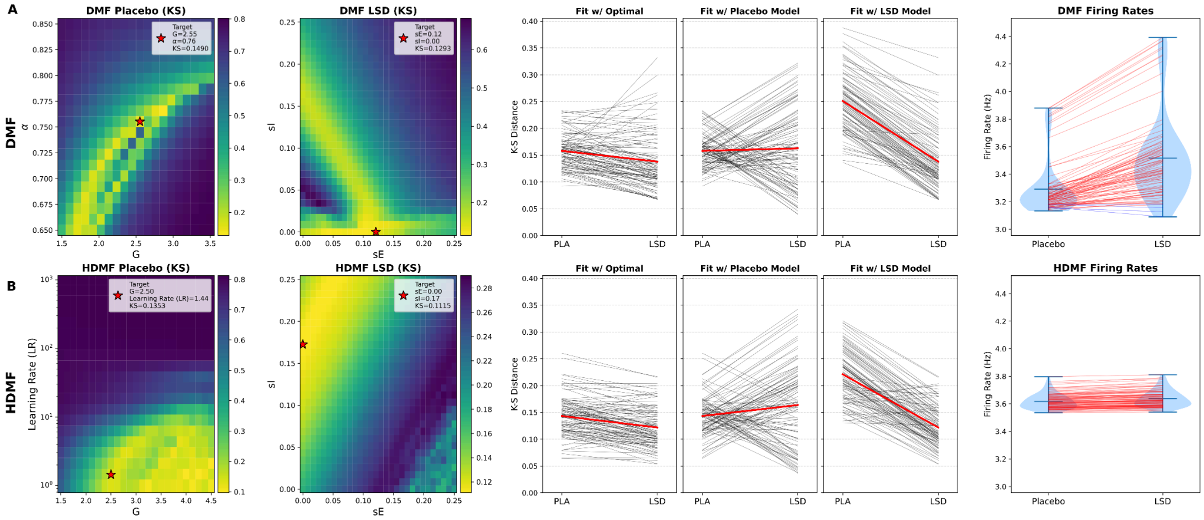


**Supplementary figure 5. Model fitting and firing-rate stability under placebo and LSD conditions.**

Comparison between the classic DMF model with static FIC and the homeostatic DMF model using the same empirical placebo/LSD dataset as in Herzog et al. Parameter optimization was performed separately for placebo and LSD by minimizing the KS distance between empirical and simulated FCD distributions; red stars indicate the selected target parameters. **A.** DMF model. Under placebo, the DMF achieved KS = 0.1490 and FC = 0.3940 at G=2.553 and α=0.755. Under LSD, after a high-resolution search at low modulation values, the optimal fit was obtained with pure excitatory modulation ($s^{(E)}$=0.119, $s^{(I)}$=0.000), improving the fit relative to placebo (KS = 0.1279, FC = 0.4211). When using 100 different random seeds, cross-fitting showed KS distances of 0.1581 ± 0.0317 for placebo and 0.1380 ± 0.0540 for LSD when using condition-specific optimal models; 0.1581 ± 0.0317 and 0.1628 ± 0.0749 when using the placebo-fitted model; and 0.2510 ± 0.0535 and 0.1380 ± 0.0540 when using the LSD-fitted model. The firing-rate panel shows increased variability and hyperexcitability under LSD. **B.** HDMF model. Under placebo, the HDMF achieved a better FCD fit than the classic DMF (KS = 0.1353, FC = 0.3448) at G=2.500 and LR=1.438. Under LSD, the HDMF reached the best global fit across the analysis (KS = 0.1115, FC = 0.3941) through pure inhibitory modulation ($s^{(E)}$=0.000, $s^{(I)}$=0.172). When using the same 100 random seeds used for the DMF, cross-fitting showed KS distances of 0.1433 ± 0.0366 for placebo and 0.1218 ± 0.0379 for LSD with condition-specific optimal models; 0.1433 ± 0.0366 and 0.1638 ± 0.0777 with the placebo-fitted model; and 0.2211 ± 0.0477 and 0.1218 ± 0.0379 with the LSD-fitted model. The HDMF maintained firing rates within a narrower range across conditions, confirming the enhanced stability under neuromodulation and enriched dynamics. Results show the mean ± standard deviation across the same 100 seeds.

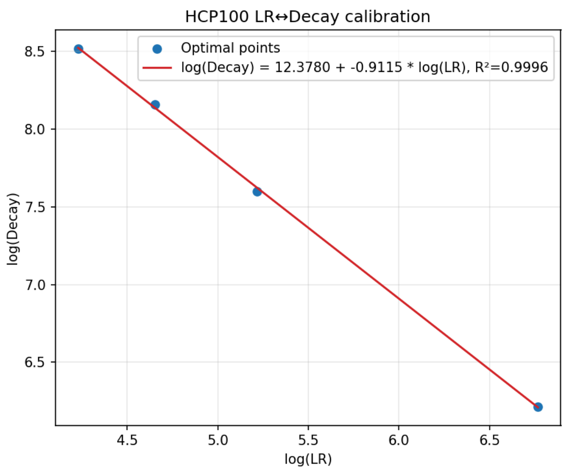


**Supplementary figure 6: Calibration of the LR-$\tau_{decay}$ relationship for the Schaefer-100 structural connectome derived from HCP data.** To avoid a full two-dimensional grid search, we selected four $\tau_{decay}$ values and, for each of them, explored a full grid of LR values. The LR value minimizing the calibration error at each $\tau_{decay}$ value was retained as an optimal point. A linear fit in log–log space captured the resulting relationship with high accuracy, allowing the parameters of the LR-$\tau_{decay}$ scaling law to be estimated efficiently.

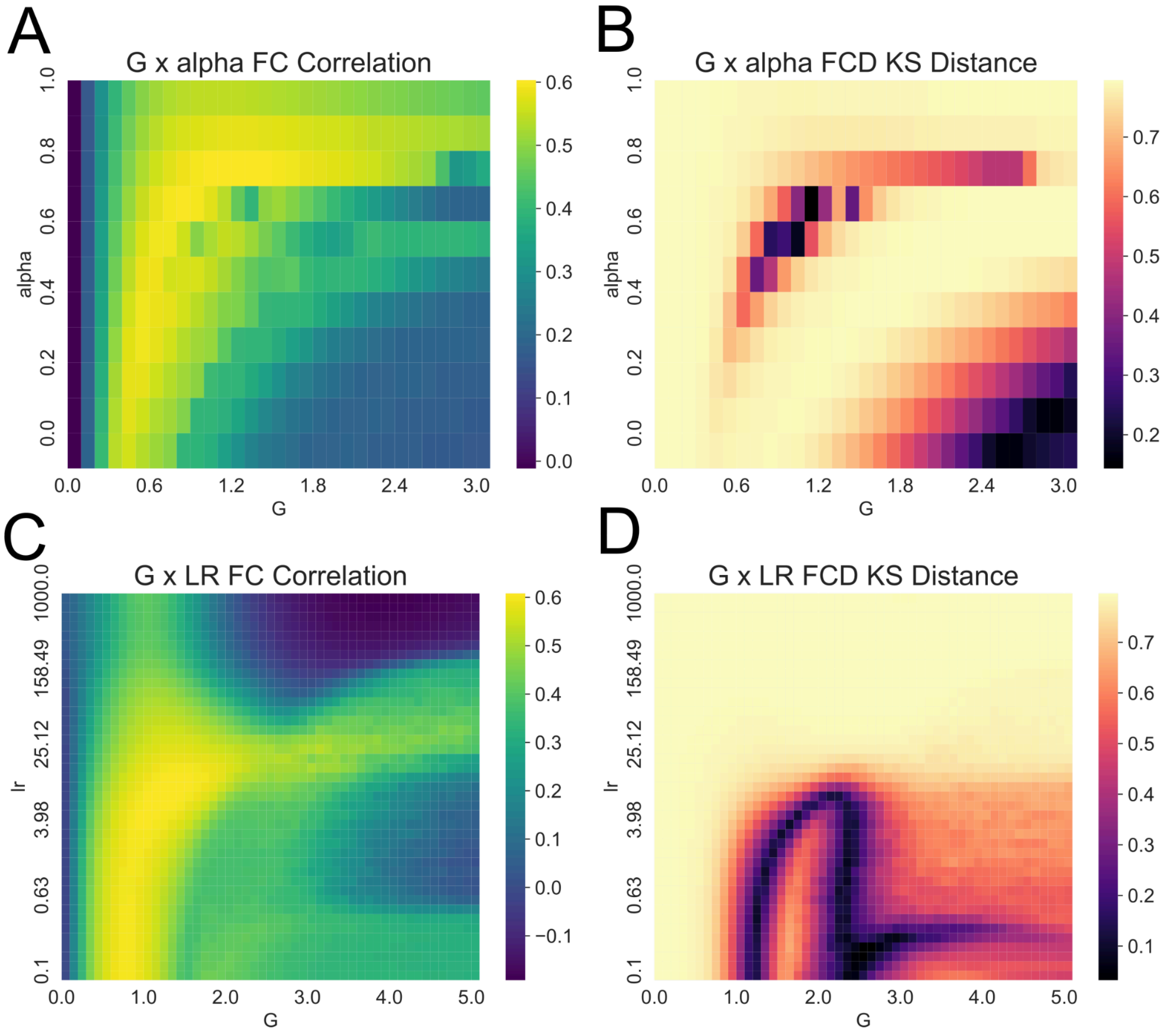


**Supplementary Figure 7. Parameter-space maps of FC and FCD fits for DMF and HDMF.** FC correlation **(A,C)** and FCD KS distance **(B,D)** shown as functions of global coupling G and α (DMF) or learning rate LR (HDMF). Regions yielding optimal FC and optimal FCD are spatially separated in parameter space for both models.

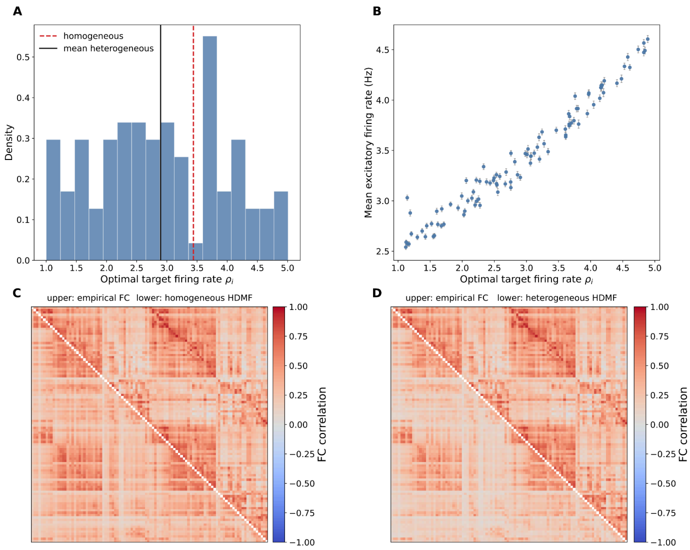


**Supplementary Figure 8. Heterogeneous target firing rates. (A)**. Distribution of the optimized regional target firing rates obtained from Bayesian optimization. Vertical lines indicate the homogeneous reference value (3.44) and the mean of the heterogeneous vector. (**B)**. Relationship between the optimized target firing rate and the mean excitatory firing rate across the same 100 seeds used for fitting; points show the across-seed mean and error bars their respective standard deviation. (**C).** Empirical FC (upper triangle) and mean simulated FC from the homogeneous HDMF model (lower triangle). (**D)** Empirical FC (upper triangle) and mean simulated FC from the heterogeneous HDMF model fitted with region-specific target firing rates.
